# High-throughput, temporal and dose dependent, effect of vitamins and minerals on chondrogenesis

**DOI:** 10.1101/811877

**Authors:** James E. Dennis, Taylor Splawn, Thomas J. Kean

## Abstract

Tissue engineered hyaline cartilage is plagued by poor mechanical properties largely due to inadequate type II collagen expression. Of note, commonly used defined chondrogenic media lack 14 vitamins and minerals, some of which are implicated in chondrogenesis. Type II collagen promoter-driven *Gaussia* luciferase was transfected into ATDC5 cells to create a chondrogenic cell with a secreted-reporter. The reporter cells were used in an aggregate-based chondrogenic culture model to develop a high-throughput analytic platform. This high-throughput platform was used to assess the effect of vitamins and minerals, alone and in combination with TGFβ1, on type II collagen expression. Significant combinatorial effects between vitamins, minerals and TGFβ1 in terms of type II collagen expression and metabolism were discovered. An ‘optimal’ continual supplement of copper and vitamin K in the presence of TGFβ1 gave a 2.5-fold increase in collagen expression over TGFβ1 supplemented media alone.

**Summary:** Current defined chondrogenic culture media lack several vitamins and minerals. Type II collagen is the quintessential marker of articular hyaline cartilage, and is commonly deficient in engineered tissue. A type II collagen promoter driven secreted luciferase construct has been transduced into ATDC5 cells and used to assess vitamin and mineral effects on chondrogenesis in a high-throughput format.

## Introduction

Osteoarthritis is a leading cause of disability with annual USA healthcare costs in excess of $500 billion (Centers for Disease Control and Prevention, 2009; Helmick et al., 2008). This laboratory has investigated media formulations and expansion surfaces to optimize cartilage formation for multiple applications (Dennis et al., 2018; Henderson et al., 2010; Henderson et al., 2007; Kean and Dennis, 2015; Kean et al., 2016a; Weidenbecher et al., 2007; Weidenbecher et al., 2008). Supplements include BMP (Mounts et al., 2012) and thyroxine (Whitney et al., 2018); nevertheless, engineered cartilage invariably contained type II collagen at levels far lower than native (Whitney et al., 2014), as is the scourge of the field. Current methods used to optimize medium conditions are tedious and inefficient. The goal of this study was to streamline the method to screen for optimal culture conditions using a high-throughput assay of a *Gaussia* luciferase reporter system linked to a type II collagen promoter. This initial study was applied to the optimization of the base medium components used to differentiate chondrocytes, using ATDC5 cells as a model system.

Many researchers, including ourselves (Dennis et al., 2017; Kean and Dennis, 2015; Kean et al., 2016a; Whitney et al., 2017), use serum-free chondrogenic media to engineer cartilage *in vitro* (Johnstone et al., 1998). Upon examination of the composition of this medium, it was noted that several vitamins and minerals potentially relevant to chondrogenesis were absent (Table 1). To address this disparity and examine their potential effects on chondrogenesis, a high-throughput assay was developed based on a *COL2A1*-*Gaussia* luciferase reporter system using ATDC5 cells formatted as aggregates in 96-well plates. The dose responses of each vitamin and mineral was assessed over time. TGFβ1 was used as a positive control, as it has previously been reported to upregulate chondrogenesis in ATDC5s (Han et al., 2005) and other mammalian cells (Puetzer et al., 2010). Combinations of vitamins and minerals which showed positive increases in type II collagen were tested in the presence of TGFβ1. This article shows proof-of-concept data for a high-throughput screen and establishes an improved chondrogenic medium for ATDC5 cells.

**Table 1.** Vitamins and minerals absent in DMEM (and therefore defined chondrogenic media (Kean and Dennis, 2015))

|  |  |  |  |
| --- | --- | --- | --- |
| **Component** | **Role in Chondrogenesis** | **Normal range<sup>1</sup>** | **References** |
| Chromium | Unknown, has a role in insulin/glucose metabolism | 0.3-28 µg/L | (Finch, 2015) |
| Cobalt | Unknown, required for Vitamin B12 and other enzymes | 0.0-0.9 µg/L | (MayoClinic; Mobasheri et al., 2006) |
| Copper | Part of lysyl oxidase complex - collagen cross-linking | 90-670 µg/L | (Finch, 2015; Makris et al., 2013a) |
| Iodine | Deficiency causes cartilage defects | 40-92 µg/L | (MayoClinic; Ren et al., 2007) |
| Manganese | Deficiency results in skeletal abnormalities | 0.6-12 µg/L | (Finch, 2015) |
| Molybdenum | Unknown | 0.3-2.0 µg/L | (MayoClinic; Sardesai, 1993) |
| α-linolenic acid | Unknown, precursor of phospholipid membrane. | 2.8-54 mg/L | (Connor, 1999; MayoClinic) |
| Thyroxine<sup>2</sup> | Enhances and stimulates terminal differentiation | 9-17 ng/L | (MayoClinic; Mello and Tuan, 2006; Whitney et al., 2017) |
| Vitamin A | Shown to inhibit chondrogenesis | 113-780 µg/L | (Lind et al., 2013; Masuda et al., 2015; MayoClinic; Pacifici et al., 1980) |
| Vitamin B7 | Unknown | 57-3004 ng/L | (MayoClinic; Ren et al., 2007) |
| Vitamin B12 | Unknown, deficiency causes growth retardation | 180-914 ng/L | (MayoClinic; Stabler, 2013) |
| Vitamin D | Implicated in chondrogenesis | 24-86 ng/L | (MayoClinic; Tsonis, 1991) |
| Vitamin E | Unknown | 3.8-18.4 mg/L | (MayoClinic; Wluka et al., 2002) |
| Vitamin K | Possible role in early chondrogenesis. | 0.1-2.2 µg/L | (Barone et al., 1991; Booth, 1997; MayoClinic) |
| Zinc | Stimulates chondrocyte growth and collagen production. | 0.6-1.2 mg/L | (Litchfield et al., 1998) (Koyano et al., 1996) |

<sup>1</sup>Normal reference range in blood or serum; synovial fluid does not appear to have been assessed for these components. Defined chondrogenic media is supplemented with vitamin C and ITS+ Premix which contains selenium and linoleic acid, thus providing 3 essential vitamins and minerals absent in DMEM. <sup>2</sup>Thyroxine, although a growth factor, was included because of our previous study showing upregulation of type II collagen (Whitney et al., 2018).

Somewhat surprisingly, the common cell culture medium Dulbecco’s Modified Eagle’s Medium (DMEM) used in cell culture of chondrocytes (and other cells) lacks several vitamins and minerals that are defined as essential (See supplemental table 1 for common media comparison). It is likely that some or all of these essential components are provided by the supplementation of fetal bovine serum during expansion, but these components are clearly absent during chondrogenic culture using defined medium (Johnstone et al., 1998; Kean and Dennis, 2015). Chondrogenic media compositions have evolved to defined media from that originally reported by Ahrens *et al*. (Ahrens et al., 1977) in 1977 to a reduced serum medium described by Ballock *et al*. (Ballock and Reddi, 1994), then to a serum-free defined medium published by Johnstone *et al.* (Johnstone et al., 1998); our current chondrogenic medium is an adaptation of the last (Kean et al., 2016b). Table 1 shows the essential vitamins and minerals lacking in DMEM, a base medium commonly used in chondrogenic differentiation by us and others. Indeed, Makris *et al*. found that copper supplementation of this base medium enhances collagen crosslinking and mechanical properties in tissue engineered cartilage (Makris et al., 2013b). To investigate this and other media supplements on chondrogenic differentiation, we assayed for *Gaussia* luciferase levels in the conditioned medium of ATDC5 cells, a commonly used chondrocyte cell line derived from mouse teratocarcinoma AT805, transduced with lentiviral *Gaussia* luciferase under the control of the *COL2A1* promoter (col2gLuc; HPRM22364-LvPG02, Genecopia). The use of the secreted form of luciferase allows for the temporal assessment of type II collagen expression in the conditioned medium of aggregate cultures.

## Results

### Pseudolentiviral particle production, infection and selection

Pseudolentiviral particles of control (eGFP) and col2gLuc were successfully produced and used to infect ATDC5 cells. Dilution titration of infected cells with eGFP determined a multiplicity of infection (MOI) of 29. Calculation of infectious particle concentration by qPCR determined this concentration to be approximately the same for col2gLuc in terms of viral particles per μl (1.9×x10^7^ and 1.6 ×10^7^ for eGFP and col2gLuc respectively), this equates to an approximate MOI of 25 for the col2gLuc studies.

### Basal Culture Optimization

In initial experiments, chondrocytes were plated in high-density pellet cultures at 50,000 and 100,000 cells/pellet at both 20% oxygen and 5% oxygen (physioxia (Kean and Dennis, 2015; Kean et al., 2016a)), with similar *Gaussia* luciferase responses to TGFβ1 in each (Fig. 1).

**Figure 1.**
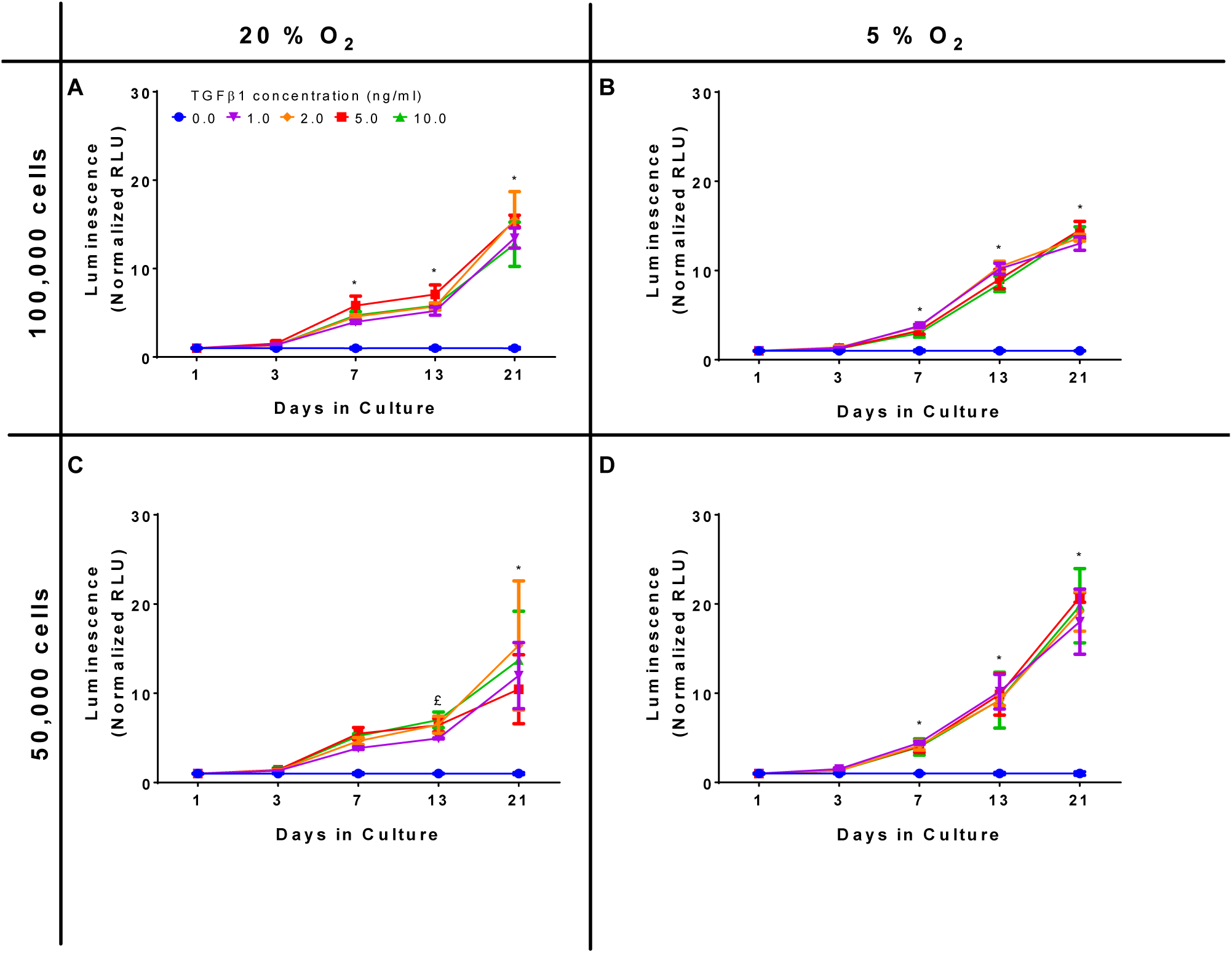
Oxygen and cell density response. Dose response curves, normalized to the control of 0 ng/ml TGFβ1 in chondrogenic media, were constructed and show a time dependent increase in luminescence but with similar responses in terms of both cell number and oxygen tension. * all concentrations significantly greater than basal media, £ concentrations ≥ ng/ml significantly greater than basal media (*p* < 0.05)

Data were averaged and the z-factor calculated according to the formula described by Zhang *et al*. (Zhang et al., 1999) 

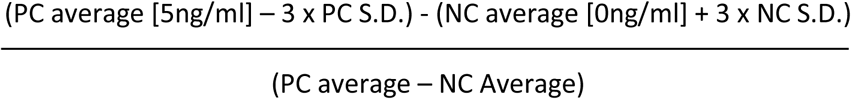

Where PC = positive control and NC = negative control

For day 21 the z’ factor for a 50,000 cell pellet was 0.89, which is regarded as very good, validating the assay as having sufficient sensitivity for high-throughput applications. Luminescent decay was assessed over 50 minutes and the half-life of positive (10 ng/ml) and negative (0 ng/ml) samples were similar; 27.8 and 33.1 minutes respectively. As significant signal to noise was achieved using a lower number of cells, the remaining experiments were all conducted using 50,000 cell aggregates in physioxia (5% O_2_) and luminescence read within 10 minutes of adding stabilized substrate.

### TGFβ1 Dose-Response

The dose-response curve of TGFβ1, a known inducer of chondrogenesis, was determined along with the 50% effective concentrations (EC50) calculated as 18, 21 and 22 pg/ml for days 7, 15 and 21, respectively (Fig. 2).

**Figure 2.**
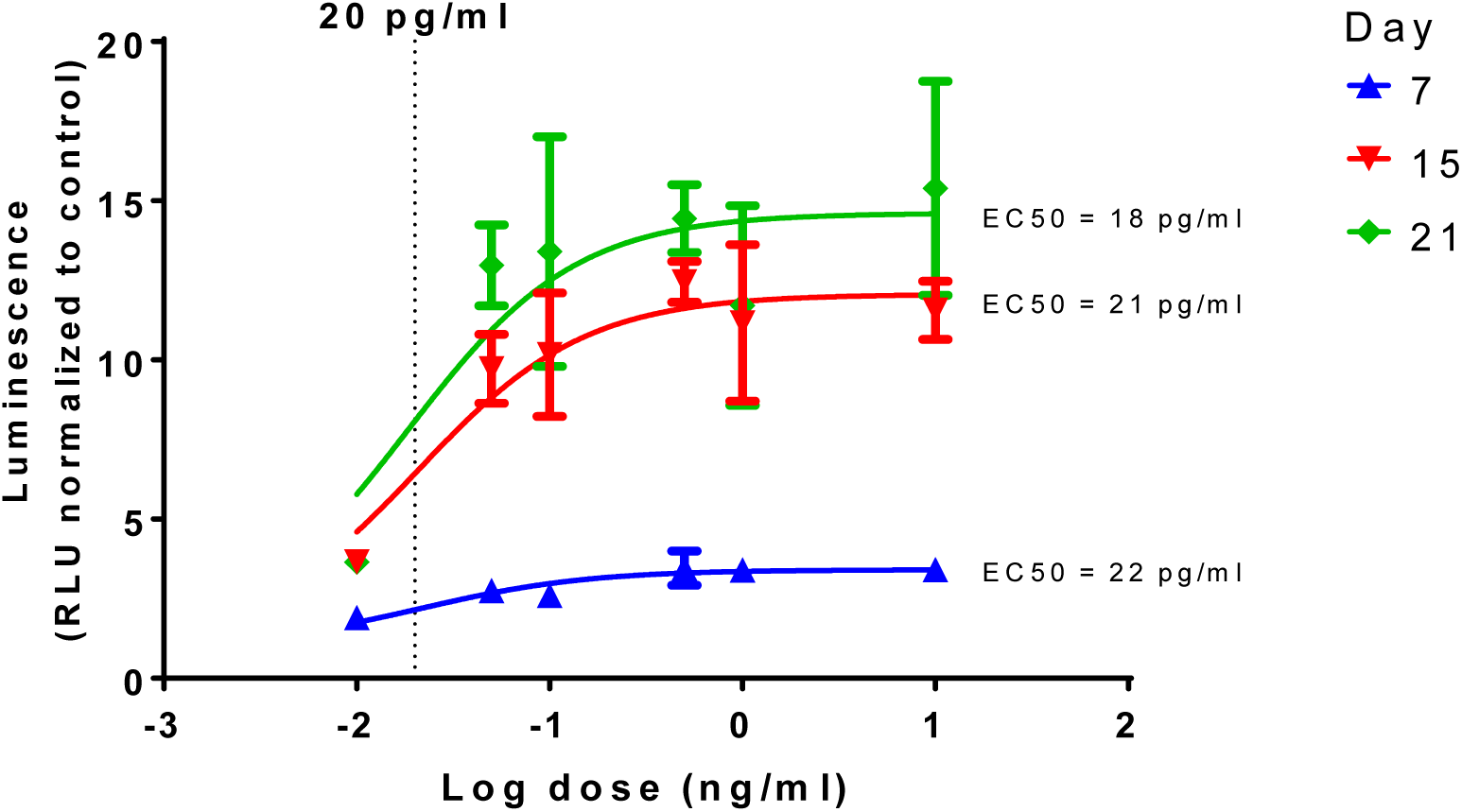
Time and dose dependent effect of TGFβ1 on chondrogenesis. Dose response curves, normalized to the control of 0 ng/ml in chondrogenic media, were constructed and show a time dependent increase in luminescence but with similar EC50s, a dotted line is shown to show 20 pg/ml in the log scale.

### Vitamin and Mineral Dose-Response in Basal Media

Vitamins and minerals were assessed at varying concentrations based on normal serum concentrations with amounts ranging from 0.01-1000 % of the maximal serum concentration. Both dose- and time-dependent effects on type II collagen expression were evident (Table 2 and Supplemental Data 1). Some additives resulted in steady expression levels over time, some with minimal effects, such as copper, manganese and molybdenum, while others showed relatively consistent, but modest, levels of upregulation, such as iodine and zinc. Vitamins B7, B12, E and K showed greater levels of upregulation that were, again, predominantly steady in nature. Vitamin A was interesting in that it showed high levels of upregulation up to day 15, but then the expression levels dropped precipitously on day 21. A similar pattern emerged for cobalt where the levels were similar to that of the basal medium control up to day 15 but dropped to approximately half those levels on day 21.

**Table 2.** Temporal effect of vitamins and minerals on chondrogenesis in basal media.

|  |  |  |  |  |  |
| --- | --- | --- | --- | --- | --- |
| **Component** | **Dose (% of maximal serum concentration<sup>†</sup>)** | **Fold change vs. basal medium** |  |  |  |
|  |  | **Week** |  |  |  |
|  |  | **0.4** | **1** | **2** | **3** |
| Chromium | 2.8 µg/L (10) | *1.73 | *2.05 | *2.29 | *2.08 |
| Cobalt | 9.00 µg/L (1000) | 1.17 | 1.45 | *4.65 | 1.03 |
| Copper | 6.70 mg/L (1000) | 0.78 | 0.68 | *8.61 | *0.16 |
| Iodine | 920 µg/L (1000) | *1.19 | 1.37 | *4.79 | 0.95 |
| Manganese | 60 µg/L (500) | 1.06 | 1.04 | *4.33 | 0.99 |
| Molybdenum | 20 µg/L (1000) | 1.25 | 1.34 | *4.69 | 1.11 |
| α-linolenic acid | 5.4 µg/L (0.1) | *1.51 | *1.37 | 1.02 | 0.82 |
| Vitamin A | 0.078 µg/L (0.01) | *1.94 | *2.22 | *3.23 | 1.06 |
| Vitamin B7 | 15.02 µg/L (500) | *1.63 | *1.68 | 1.22 | 1.36 |
| Vitamin B12 | 91.4 ng/L (10) | *1.68 | *1.70 | *1.63 | 1.37 |
| Vitamin D | 8.6 ng/L (10) | *1.31 | 1.02 | 0.86 | *0.68 |
| Vitamin E | 18.4 mg/L (100) | *1.51 | *1.56 | *1.44 | 1.16 |
| Vitamin K | 22 µg/L (1000) | 0.97 | 0.94 | *5.68 | 1.04 |
| Zinc | 6.0 mg/L (500) | 1.27 | 1.54 | 1.10 | *1.51 |

<sup>†</sup>see Table 1 for normal reference range in blood or serum; synovial fluid does not appear to have been assessed for these components; defined chondrogenic media is supplemented with vitamin C and ITS+ Premix which contains selenium and linoleic acid, which are absent in DMEM. \* Significantly different vs. basal medium, 2-way ANOVA $p < 0.05$ Sidak's multiple comparison test.

### Vitamin and Mineral Combinatorial Effects

A selection of those factors that had overall positive effects on the relative luminescence of the conditioned media (vitamin B7, copper, iodine, thyroxine and zinc) were tested at their optimal individual concentration in basal medium and in combination with TGFβ1 (1 ng/ml). These 6 factors gave 64 conditions; 53 of which were discovered to be significantly different vs. the basal medium, with 46 being increased and 7 being decreased at day 21 (Supplemental Data 2). An abbreviated selection of conditions in the presence of TGFβ1 is shown in Figure 3 (18 of 64); several conditions in combination with TGFβ1 are greater than TGFβ1-supplemented basal medium alone (15 of 31; Fig. 3 and Supplemental Data 2). The results showed several interesting temporal patterns.

**Figure 3.**
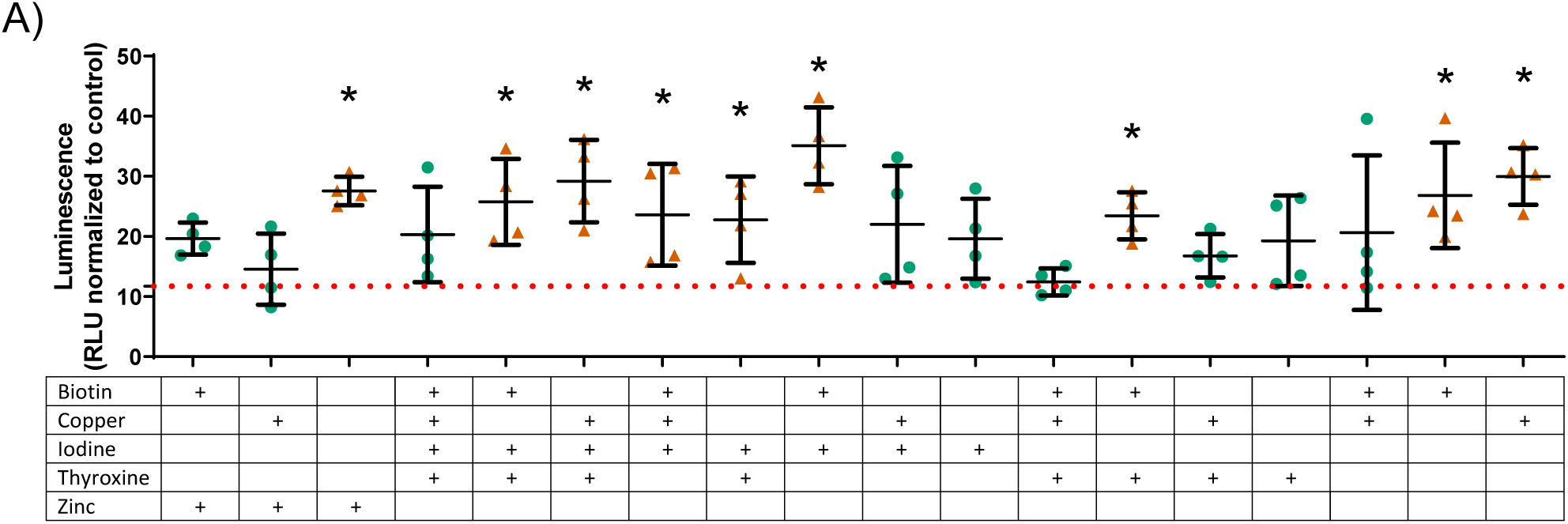
Collagen expression with various vitamin and mineral combinations in the presence of TGFβ1 on day 21. Vitamins and minerals which had been identified as promoting type II collagen expression in the initial screen were tested in combination at their individually optimized concentration in the presence of TGFβ1 at 1ng/ml. The values are normalized to the plate control (basal media). Green circles (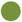) indicate values that are significantly greater than those for basal media alone, and the dashed red line indicates the luminescence achieved with TGFβ1 supplementation at 1 ng/ml. The orange triangle (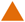) and an * indicate values that were discovered different, with false discovery rate set at 1%.

Due to the prohibitively large number of combinations with the remaining 10 factors (1023 conditions), the first combination experiments (designated here as supplemented media 1-4) were additionally supplemented with combinations of vitamin B12, vitamin E, vitamin K and chromium, each of which had been determined to effect an increase in relative luminescence. The individually determined optimal concentrations in the 3 highest performing combinations from the previous experiment were used for this phase of the study. The TGFβ1 supplemented basal media is termed Sup 1; Sup 2 = Sup 1 + iodine + biotin; Sup 3 = Sup 1 + copper; Sup 4 = Sup 1 + iodine + thyroxine + copper. This resulted in assessment of an additional 60 conditions. In this set of experiments, Sup 3 was higher than Sup 1, as was shown previously in Figure 3, and several combinations were lower than the supplemented media in this and Sup 4 comparisons (Fig. 4 and Supplemental Data 3). Although all combinations significantly increased type II collagen expression over that of basal medium, only vitamin K significantly increased luminescence above TGFβ1- and copper-supplemented basal medium (2.5-fold vs TGFβ1 alone; Sup 3; Fig. 4C and Supplemental Data 3). However, several combinations showed luminescence levels lower than baseline supplemented media (Sup 3 and Sup 4).

**Figure 4.**
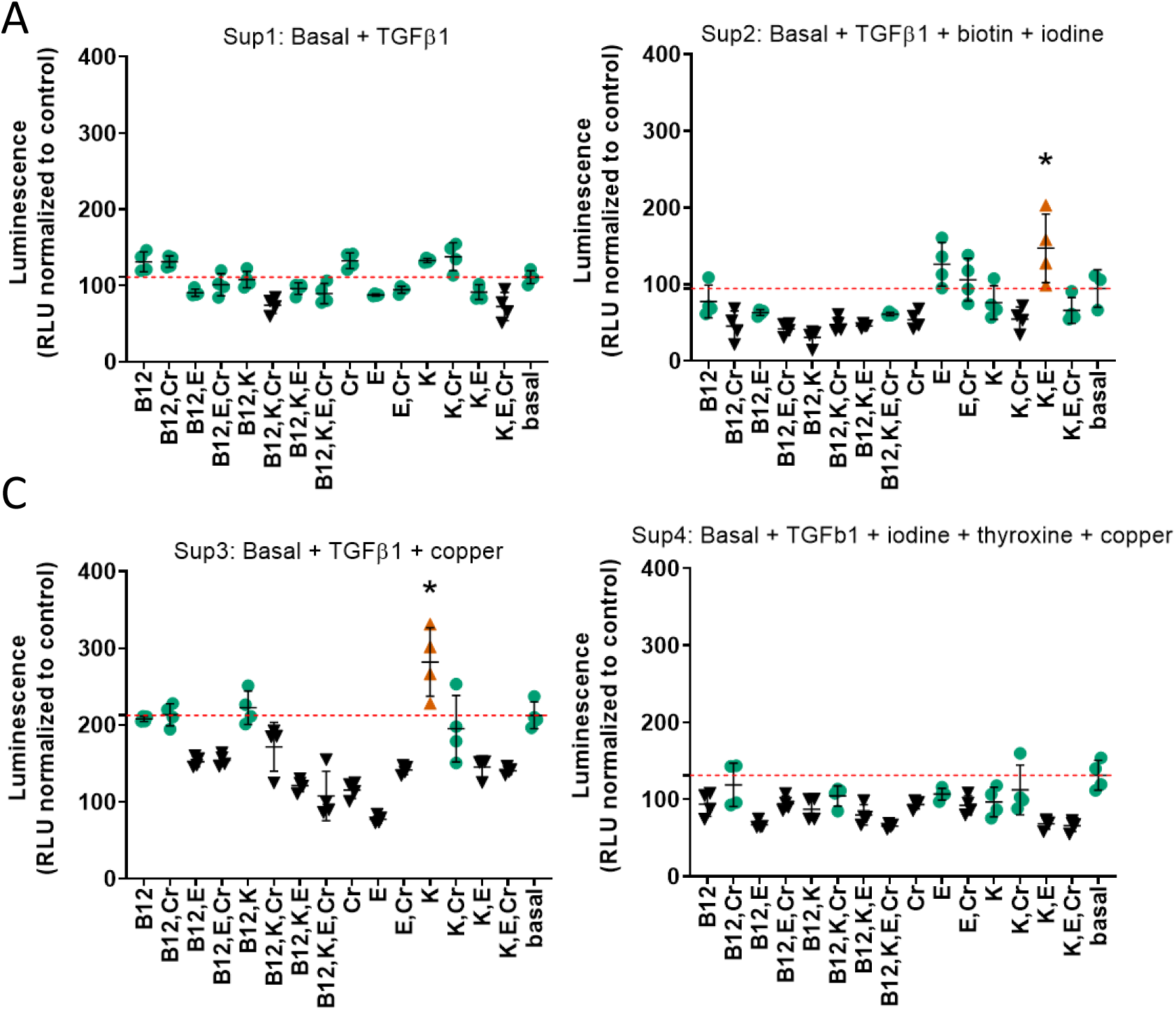
Combinations of chromium, vitamins B12, E and K in supplemented media at day 21. Vitamins and minerals which had been identified as promoting type II collagen expression in the initial screen were tested alone and in combination in supplemented medias 1-4 at their individually optimized concentration in basal media. The values are normalized to the plate control (basal media); the red dashed red line indicates the luminescence achieved with that particular supplemented media alone. Conditions that were statistically greater than their respective supplemented media are represented by a 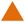 and an *, black symbols ▾ indicate that those values were lower than the respective supplemented media. n ≥ 3 ± S.D.

### Dose response to vitamin and mineral supplementation in the presence of TGFβ1

Given the complex responses seen with combinations in the presence of TGFβ1, vitamin and mineral dose responses (0.01-500 % serum max) were further tested in the presence of TGFβ1 (1 ng/ml). Both time and dose responses were evident (Table 3, Fig. 5 and Supplemental Data 4). Conditioned medium was collected on days 3, 7, 15 and 21 and assayed for *Gaussia* luciferase expression to evaluate the temporal effects of different additives. The results were expressed as the fold-change relative to time-matched basal medium controls (Table 3).

**Table 3.** Dose and temporal effect of vitamins and minerals in the presence of TGFβ1 on chondrogenesis.

|  |  |  |  |  |  |
| --- | --- | --- | --- | --- | --- |
| **Component** | **Dose (% of maximal serum concentration<sup>‡</sup>)** | **Fold change vs. basal medium** |  |  |  |
|  |  | **Day** |  |  |  |
|  |  | **3** | **7** | **15** | **21** |
| Chromium | 28 ng/L (0.1)* | 2.2 | 8.2 | 29.1 | 43.3 |
| Cobalt | 90 pg/L (0.01)* | 1.8 | 5.0 | 22.1 | 39.5 |
| Copper | 67 ng/L (0.01)* | 1.9 | 5.6 | 23.5 | 41.3 |
| Iodine | 92 ng/L (0.1) | 2.5 | 5.4 | 17.4 | 25.4 |
| Manganese | 12 ng/L (0.1) | 1.8 | 5.8 | 16.4 | 21.6 |
| Molybdenum | 2 ng/L (0.1) | 1.7 | 5.2 | 15.3 | 20.5 |
| α-linolenic acid | 27 mg/L (50) | 2.0 | 7.1 | 27.1 | 34.8 |
| Thyroxine | 25 ng/L (147)* | 1.7 | 6.0 | 18.1 | 22.6 |
| Vitamin A | 78 ng/L (0.01) | 2.3 | 5.5 | 15.7 | 24.8 |
| Vitamin B7 | 3 ng/L (0.1) | 1.6 | 5.0 | 15.5 | 21.4 |
| Vitamin B12 | 914 ng/L (100)* | 1.9 | 7.2 | 35.5 | 52.2 |
| Vitamin D | 86 pg/L (0.01)* | 1.7 | 5.1 | 20.0 | 39.2 |
| Vitamin E | 18.4 mg/L (100) | 1.6 | 6.6 | 21.7 | 27.1 |
| Vitamin K | 11.0 µg/L (500)* | 2.6 | 8.1 | 24.7 | 36.2 |
| Zinc | 0.12 µg/L (0.01)* | 1.7 | 5.0 | 19.0 | 37.2 |
| TGFβ1 medium | 1 µg/L (956<sup>†</sup>) | 1.7 | 5.3 | 16.3 | 28.1 |

<sup>‡</sup>see table 1 for normal reference range in blood or serum. <sup>†</sup> Maximum normal adult serum TGFβ1 is 104.6 pg/ml (Tas et al., 2014), this experiment used 1 ng/ml. No data is available for synovial fluid; defined chondrogenic media is supplemented with vitamin C and ITS+ Premix which contains selenium and linoleic acid, which are absent in DMEM. \* Slope or intercept of linear regression significantly different compared to TGFβ1 supplemented basal medium $p < 0.05$ .

**Figure 5.**
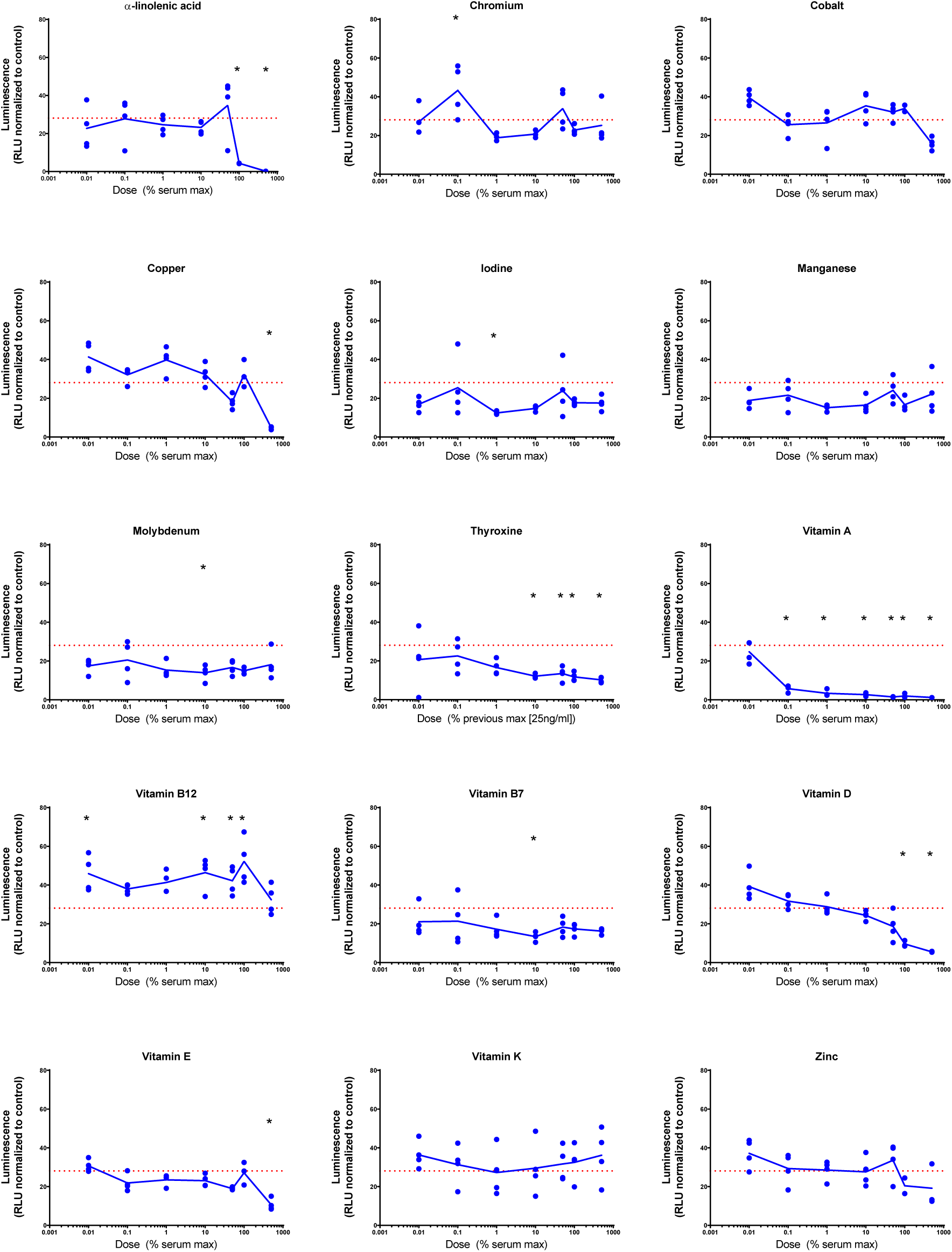
Day 21 evaluation of collagen expression in vitamin and mineral supplemented TGFβ1 supplemented basal medium. Conditioned media was sampled at day 21 and assessed for luminescence, and normalized against the plate control basal medium. The red dashed line indicates the luminescence of the TGFβ1 supplemented basal medium. * Significantly different vs. TGFβ1 supplemented basal medium, 2-way ANOVA *p* < 0.05 Sidak’s multiple comparison test.

Summation of the fold-change across the whole experiment to give area under the curve analysis identified 40 conditions that were greater than the TGFβ1 supplemented medium. The top 10 conditions that gave greater expression over the whole experiment are shown in Table 4 with an example plot in Fig. 6 (the remaining comparisons are shown in Supplemental Data 6).

**Table 4.**
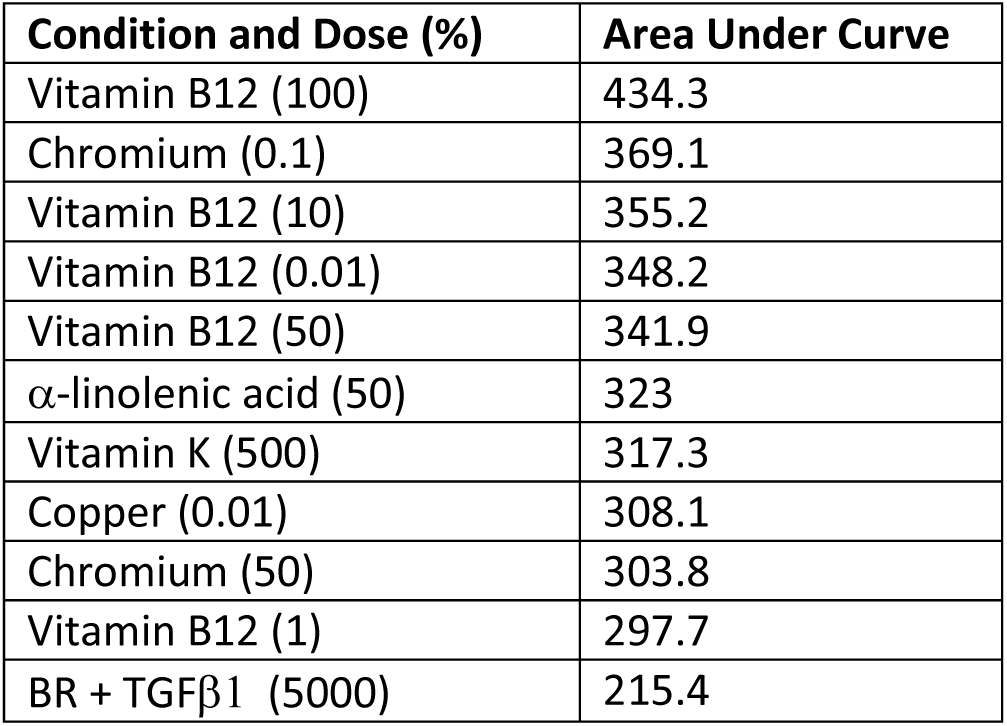
Overall dose and time response.

**Figure 6.**
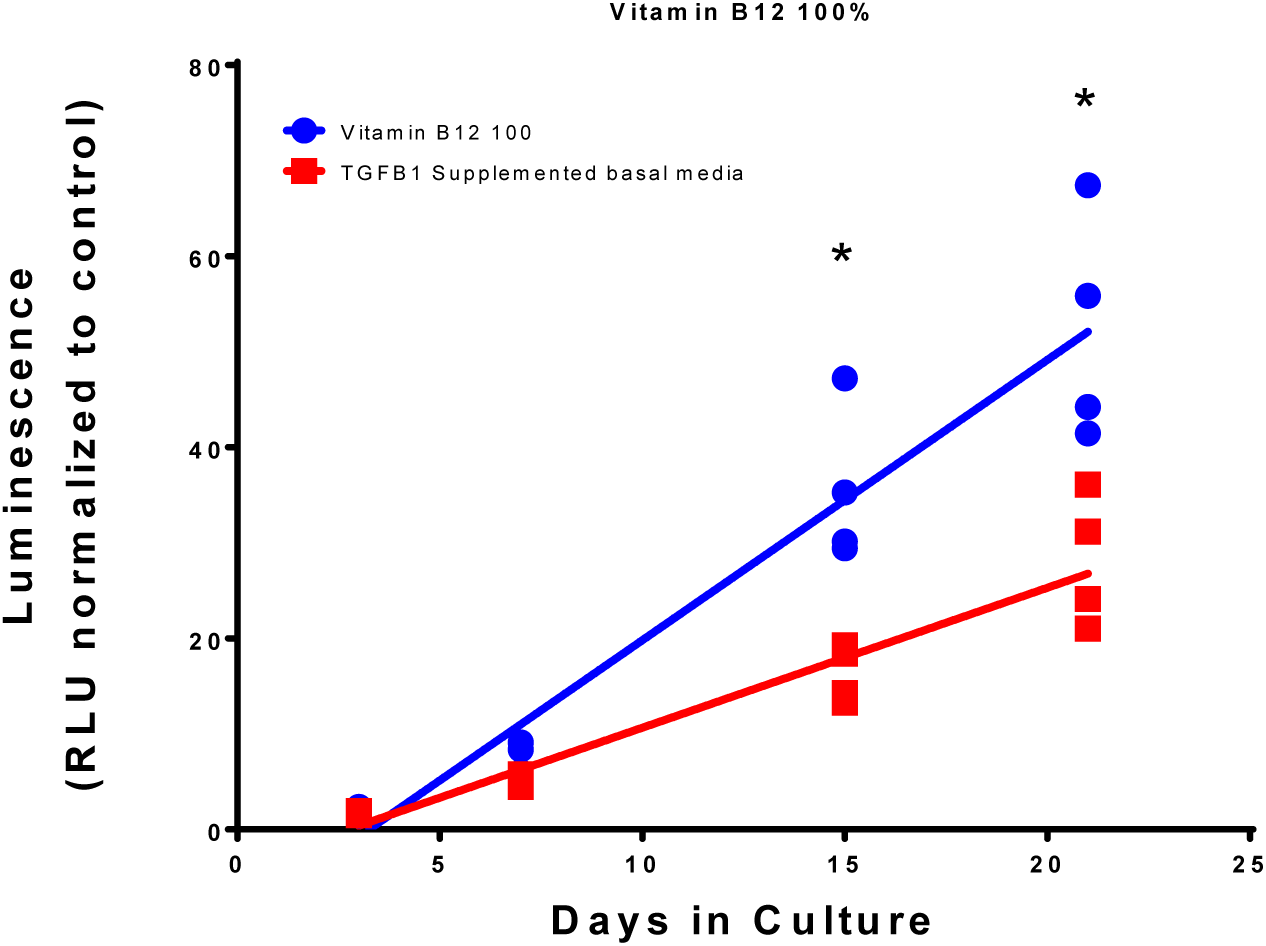
Comparison of Vitamin B12 and TGFβ1 supplemented medium with TGFβ1 supplemented basal medium. Type II collagen driven luminescence was assessed from the conditioned media and normalized to the plate basal medium control. Lines indicate a linear regression; symbols indicate individual data points. * Significant difference vs. TGFβ1 supplemented basal medium; 2-way ANOVA p < 0.05 Sidak’s multiple comparison test.

To assess the effects of supplementation on metabolic activity, resazurin assays were performed at day 21. At day 21, increased metabolic activity was detected at low doses of cobalt, copper, vitamin D, vitamin E and zinc (Fig. 7 and Supplemental Data 5). Decreased metabolic activity was seen at high doses of copper and vitamin D. Vitamin A was predominantly inhibitory and vitamin B12 predominantly stimulatory. When type II collagen-driven luciferase expression was normalized against resazurin metabolism, none of the components tested showed increased expression at day 21 compared to control. Indicating that, at this time point, the effect was predominantly or exclusively linked to metabolic activity.

**Figure 7.**
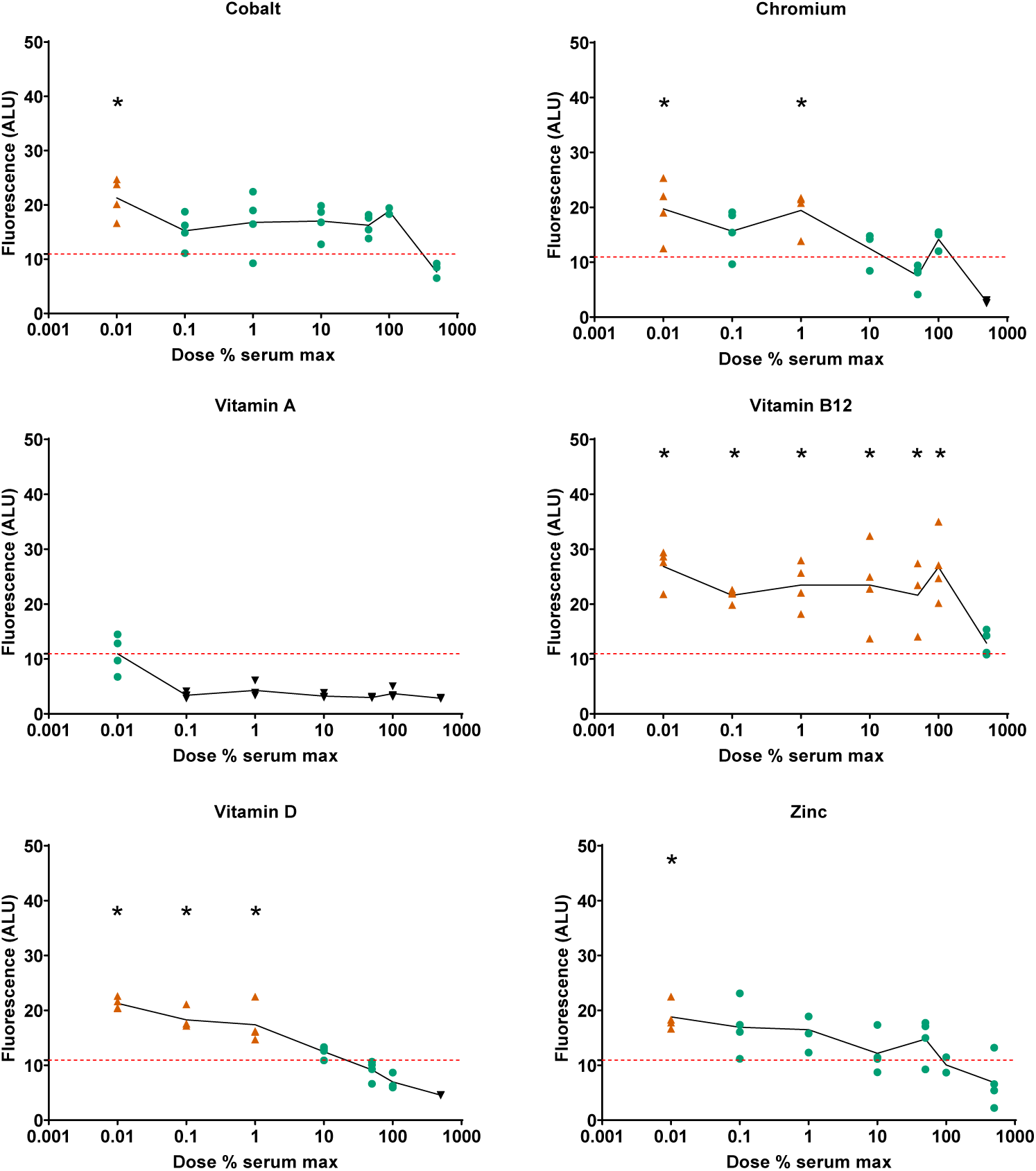
Effect of vitamin and mineral addition to TGFβ1 supplemented medium on metabolic activity. Cells were treated with alamar blue, media was sampled at day 21 and assessed for fluorescence. The red dashed line indicates the fluorescence of the TGFβ1 supplemented basal medium. An orange triangle (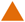) and an * indicate greater than TGFβ1 supplemented basal medium; black triangles (▾) indicate lower vs. TGFβ1 supplemented basal medium, 1-way ANOVA *p* < 0.05 Dunnett’s multiple comparison test.

## Materials and Methods

### ATDC5 cell culture and transformation

ATDC5 cells (a kind gift of Dr. Florent Elefteriou, Baylor College of Medicine) were grown in DMEM-LG (Hyclone) containing 5% FBS (Atlanta Biologicals) and 1% penicillin/streptomycin (GIBCO) at 37 °C in a humidified atmosphere with 20% O_2_, 5% CO_2_. Cells were transformed with pseudolentiviral particles during log expansion phase at approximately 30% confluence. Concentrated lentivirus was diluted with growth medium and mixed 1:1 with polybrene (8 μg/ml; EMD Millipore) in Opti-MEM (Gibco), then applied to the cells. This concentration of polybrene was determined to have minimal effects on proliferation and significantly improved infection. Plates were incubated at 4°C for 10 minutes before transfer to 37°C (humidified atmosphere with 20% O_2_, 5% CO_2_) for 12h. Media were then exchanged for growth medium and cells were grown to ∼90% confluence. Cells were then trypsinized and re-plated at 6,000 cells/cm^2^. After allowing cells to adhere for 24h, infected cells were selected by incubation in growth medium containing puromycin (2 μg/ml; Alfa Aesar) for 1 week. This concentration was determined to be the lowest concentration with >95% uninfected cell death on human mesenchymal stromal cells and chondrocytes (data not shown). Puromycin-selected cells were then grown in normal growth medium for no more than 4 passages before use in differentiation experiments.

### Lentiviral Construct

The collagen type II promoter (COL2A1)-driven, secreted *Gaussia* luciferase was custom ordered (GeneCopoeia; HIV-based lentiviral third generation; HPRM22364-LvPG02). The promoter length is 1,426 bp. This plasmid (col2gLuc; HPRM22364-LvPG02), the eGFP control (EX-EGFP-Lv105; GeneCopoeia) the packaging plasmid (psPAX2) and the envelope plasmid (pMD2.G) were amplified in E.coli (GCI-L3; GeneCopoeia), and purified with a silica column (Qiagen Maxiprep). Plasmids were confirmed by restriction digest and agarose gel electrophoresis. Pseudolentiviral particles were made in HEK-293Ta cells (GeneCopoeia) by co-transfection of psPAX2, pMD2.G and HPRM22364-LvPG02 using calcium phosphate precipitation (Dull et al., 1998). Pseudolentiviral particles were harvested from conditioned medium at 48h and concentrated by centrifugation (10,000 RCF, 4 °C, overnight). Infectious particle concentration was estimated by serial dilution of pseudolentiviral particles and flow cytometry assessment of eGFP positive cells produced in the same batch (Cellecta, 2018). Lentiviral particle concentration was further determined by qPCR assessment of lentiviral RNA according to the manufacturer’s instructions (Lenti-Pac titration kit; LT005; GeneCopoeia).

### Chondrogenic Differentiation

Cells were differentiated in pellet culture at 37 °C in a humidified atmosphere with 5% O_2_, 5% CO_2_ (50,000-250,000 cells/pellet) in polypropylene 96-well plates (Phenix) in defined chondrogenic medium: DMEM-HG supplemented with 1% insulin, transferrin, selenium + premix; 130 mM ascorbate-2-phosphate, 2 mM GlutaMax, 1% sodium pyruvate, 1% MEM non-essential amino acids, 100 nM dexamethasone, 1.25 μg/ml fungizone, 1% penicillin/streptomycin. This medium was further supplemented with vitamins and minerals (Table 4), thyroxine (Whitney et al., 2018) and recombinant human TGFβ1 (Peprotech). Medium sampling, mixing and exchange were conducted with a Tecan Freedom Evo fitted with a 96-head multi-channel aspirator (MCA) and 4-channel dispensing tips (DiTi). The pipetting profile for the MCA was set to aspirate at 10 μl/s and dispense at 500 μl/s. The pipetting profile for the DiTi was set to aspirate at 20 μl/s, 100 μl/s and 150 μl/s for < 15 μl, 15-200 μl and 200-1000 μl respectively and to dispense at 600 μl for all volumes.

**Table 4.** Sources and stock solution preparation of vitamins and minerals.

| Component | Format | Stock solution | Catalog # | Supplier |
| --- | --- | --- | --- | --- |
| Chromium | CrCl<sub>3</sub>·6H<sub>2</sub>O | H<sub>2</sub>O | 27096 | Sigma-Aldrich |
| Cobalt | CoCl<sub>2</sub>·6H<sub>2</sub>O | H<sub>2</sub>O | C8661 | Sigma-Aldrich |
| Copper | CuSO<sub>4</sub>·5H<sub>2</sub>O | H<sub>2</sub>O | C7631 | Sigma-Aldrich |
| Iodine | NaI | H<sub>2</sub>O | A15480 | Alfa Aesar |
| Manganese | MnCl<sub>2</sub>·4H<sub>2</sub>O | H<sub>2</sub>O | M3634 | Sigma-Aldrich |
| Molybdenum | (NH<sub>4</sub>)<sub>6</sub>Mo<sub>7</sub>O<sub>24</sub>·4H<sub>2</sub>O | H<sub>2</sub>O | M1019 | Sigma-Aldrich |
| α-linolenic acid | CH<sub>3</sub>(CH<sub>2</sub>CH=CH)<sub>3</sub>(CH<sub>2</sub>)<sub>7</sub>CO<sub>2</sub>H | Ethanol | L2376 | Sigma-Aldrich |
| Thyroxine | 3,3',5-Triiodo-L-thyronine sodium salt | 1M NaOH | T6397 | Sigma-Aldrich |
| Vitamin A | Retinoic acid | Ethanol | 44570 | Alfa Aesar |
| Vitamin B7 | D(+)-Biotin | H<sub>2</sub>O | A14207 | Alfa Aesar |
| Vitamin B12 | Cyanocobalamin | H<sub>2</sub>O | V6629 | Sigma-Aldrich |
| Vitamin D | Calcitriol | Ethanol | S1466 | Selleck Chemicals |
| Vitamin E | (+/-)-A-Tocopherol | Ethanol | T3251 | Sigma-Aldrich |
| Vitamin K | Menaquinone (K2) | Ethanol | 47774 | Sigma-Aldrich |
| Zinc | ZnSO<sub>4</sub>·7H<sub>2</sub>O | H<sub>2</sub>O | 33399 | Alfa Aesar |

### Luciferase assessment

Conditioned cell culture medium was sampled from plates (20 μl/well; MCA) at feeding and pipetted into a white 96-well plate (Greiner) and assessed for luminescence using the stabilized *Gaussia* luciferase kit (NEB; 50 μl of a 1:1 dilution with 18 MΩ water) and read in a plate reader (Tecan M200 Pro; 2s/well, no attenuation). Basal conditioned medium from cells was supplemented with vitamins and minerals to determine if there was any effect on the luciferase assay, none was seen (Supplemental Figure 7).

### Metabolic assay

Metabolic activity was assessed on day 21 using the resazurin assay (O’Brien et al., 2000; Schreiber-Brynzak et al., 2015). Resazurin (Sigma) was dissolved in phosphate buffered saline to give a 10x stock solution (500 µM), sterile filtered (0.2 µm) and aliquots made and frozen (−80 °C). Frozen aliquots were thawed and diluted in phosphate buffered saline to 50 µM, 20 µl of this solution was then added to each well and incubated at 37 °C for 6 h. Fluorescence was read (Tecan Infinite 200 Pro) with excitation at 530 nm and emission at 560 nm.

### Statistics

Statistics were performed using GraphPad Prism (GraphPad Software, Inc.). Comparisons of dose response over time were made using a two-way ANOVA with Sidak’s multiple comparison post-hoc test. Comparisons between supplemented media (Sup1-4) and additives were made using a two-way ANOVA with Sidak’s multiple comparison post-hoc test. Comparisons between basal or TGFβ1 supplemented media with combinations were made using multiple t-tests with a false discovery rate method of Benjamini and Hochberg set at 1%. Comparisons of the metabolic dose response due to supplementation with vitamins and minerals were made using ANOVA with Dunnett’s multiple comparison post-hoc test.

## Discussion

Tissue engineering of cartilage has the potential to ameliorate disability due to arthritis by a) providing a better *in vitro* model for drug discovery and b) providing a biological replacement eliminating the need for total joint arthroplasty. Without a permissive environment, i.e. one that contains all the vitamins and minerals that have been deemed essential, reflecting *in vivo* conditions, drug discovery and tissue engineering efforts are potentially hampered by inappropriate responses. This is exemplified in this study through the synergistic effect of vitamins, minerals and TGFβ1 on the expression of type II collagen (Fig 3). It is perhaps unsurprising that vitamins and minerals which are known to be essential for proper development have significant effects on collagen expression in this *in vitro* system. It is therefore surprising that they are omitted from common basal media; this work establishes that, while not all components promote type II collagen expression, they can still have an effect on chondrogenesis, such as through the stimulation or inhibition of metabolism. These complex temporal effects are to be expected given the broad range of enzymes, signalling pathways and physiological processes that require these vitamins and minerals for their activity.

Multiple laboratories have focused on methods to engineer functional cartilage for various applications. In many instances, the mechanical properties fall short of that of native cartilage tissue. One major concern is that the diminished biomechanical properties of engineered cartilage are predominantly a result of insufficient production of type II collagen, which is the most abundant molecule in cartilage after water (Sophia Fox et al., 2009). Results in our laboratory have shown type II collagen content to be approximately 20% of that in native tissue (Whitney et al., 2018). Other laboratories have reported similar results, such as those of Sato *et al*., who showed type II collagen levels from cultured human MSCS at 20% that of native tissue, even after enhancement by the addition of epigallocatechin-3-gallate, which doubled the type II collagen output (Sato et al., 2017). Similar results were shown by Gemmiti and Guldberg, where collagen content of bioreactor-grown bovine chondrocytes was 25% that of native collagen content (Gemmiti and Guldberg, 2006), and in a study by Shahin and Doran where type II collagen levels were 18% that of native tissue (Shahin and Doran, 2011). Other studies have shown collagen levels at less than 10% of native tissue (Mahmoudifar and Doran, 2010; Tian et al., 2013).

The goal of the present study was to streamline the methodology for optimizing type II collagen expression and, at the same time, using the examination of vitamins and minerals missing from conventional defined media both as a proof of concept and as a much needed inquiry. To accomplish that goal, an aggregate chondrogenesis assay (Yoo et al., 1998) that had been modified for a 96-well format (Penick et al., 2005) was applied as a means to use a high-throughput assay to screen chondrocytes transduced with a *Gaussia* luciferase reporter driven by a type II collagen promoter. This has resulted in significant insights into the role of vitamins and minerals on chondrocyte metabolism. There is a clear interaction between TGFβ1 and some vitamins and minerals (copper, vitamin A). However, even at our reduced TGFβ1 concentration of 1ng/ml, one tenth of that commonly used but still ∼500-times the EC_50_ and ∼10-times normal serum concentration, we may have masked some potential stimulators.

To date, only a few attempts at developing high-throughput assays for chondrogenesis have been reported. Greco *et al*., developed what they termed a “medium-throughput” assay that used the human C-28/12 chondrocyte cell line where cells were seeded as 1.0 × 10^4^ micromasses in 24-well plates; terminal assays were used to determine gene expression and GAG content (Greco et al., 2011). Another high-throughput assay, using only 1.0 × 10^4^ cells per well, was used to screen for factors influencing mesenchymal stem cell chondrogenesis (Huang et al., 2008) where, again, terminal assays were used to assess gene expression and GAG content. A similar aggregate assay was also described for adipose-derived stem cells where 2.0 – 5.0 × 10^5^ cells per well of a 96-well plate were cultured for several weeks and then assayed for GAG content as a terminal assay (Abu-Hakmeh and Wan, 2014). Another study used ATDC5 cells in a high-throughput assay to test a library of factors for their effect on total collagen production (Le et al., 2015). In their study, ATDC5 cells were plated at 5.0 × 10^3^ cells per well and allowed to expand for 3 days and, after 6 days, were assayed for collagen content using a fluorescent collagen-binding probe (CN35-AF488) to measure collagen content. In all of these cases, the assays were conducted as terminal assays, that is, at the end of the incubation period. One of the strengths of the methodology shown here is that the assays are conducted on conditioned medium, allowing for a complete temporal assessment of the different factors, and combinations of factors, over time.

Whilst the addition of components to media formulations makes their composition more complex, we would argue that, in order to model the effect of any *in vivo* manipulation, a permissive environment, i.e. one that contains factors essential for normal cell metabolism is necessary. Given that: 1) cells are relatively efficient in recycling many of their components, 2) *in vitro* systems are an imperfect model, and 3) the conditions that exist within a developing joint are relatively unknown, our current suggested ‘optimal’ medium for chondrogenesis is: DMEM-HG supplemented with 1% insulin, transferrin, selenium + premix; 130 mM ascorbate-2-phosphate, 2 mM GlutaMax, 1% sodium pyruvate, 1% MEM non-essential amino acids, 100 nM dexamethasone, 1.25 μg/ml fungizone, 1% penicillin/streptomycin, 1 ng/ml TGFβ1, 670 ng/ml copper, 2.2 ng/ml vitamin K (highest yield, fig. 4). However, based on the apparent shift around day 15 both in this work and that shown by Huynh *et al*. (Huynh et al., 2018) there is potential for a media switch around this time. In addition, we consider that a permissive medium solution would contain other vitamins and minerals at 1/100^th^-1/1000^th^ of their serum max, potentially allowing somewhat of a ‘normal’ response to other manipulations such as mechanical stimulation (Nazempour et al., 2016). The proposed supplements to chondrogenic medium are shown in Table 5 for the two time periods 0-15 and 16-21 days. Select vitamins and minerals were removed from the later days in culture as they decreased collagen expression after 15 days; this may be because the tissue is more mature, and they are recycled more efficiently and/or used less. The fat soluble vitamins A, D, E, K and α-linolenic acid should be combined with ITS+ premix as a stock solution as they will bind to the bovine serum albumin in that solution. These suggested media need further investigation in this and different systems.

**Table 5.** Proposed supplements to defined chondrogenic media.

| Component | Final media concentration | Day 0-15 | Day 16-21 |
| --- | --- | --- | --- |
| α-linolenic acid | 5.4 mg/L | ✓ | ✓ |
| Chromium | 28 ng/L | ✓ | ✓ |
| Cobalt | 90 pg/L | ✓ | ✓ |
| Copper | 6.7 μg/L | ✓ | ✓ |
| Iodine | 92 ng/L | ✓ | ✓ |
| Manganese | 1.2 ng/L | ✓ | ✗ |
| Molybdenum | 2.0 ng/L | ✓ | ✗ |
| Thyroxine | 25 ng/L | ✓ | ✗ |
| Vitamin A | 78 ng/L | ✓ | ✗ |
| Vitamin B7 | 1.5 μg/L | ✓ | ✗ |
| Vitamin B12 | 914 ng/L | ✓ | ✓ |
| Vitamin D | 86 pg/L | ✓ | ✓ |
| Vitamin E | 18.4 mg/L | ✓ | ✓ |
| Vitamin K | 1.1 μg/L | ✓ | ✓ |
| Zinc | 120 ng/L | ✓ | ✓ |

## Conclusions

A high-throughput amenable, temporal, chondrogenesis assay has been developed in a model chondrocyte cell line, ATDC5. Medium supplementation with vitamins and minerals resulted in changes in type II collagen expression and metabolic activity dependent on both time and concentration.

## Acknowledgements

The authors are extremely grateful to Dr. Ken Scott (1974-2017) for his permitting us the use of his Tecan Evo Freedom, without which we could not have pursued this research. We would also like to acknowledge Dr. D. MacKay for proofreading and funding from the Baylor College of Medicine (J.E.D.) and the Bone Disease Program of Texas (T.J.K.).

