## Supplemental Data for "High-throughput, temporal and dose dependent, effect of vitamins and minerals on chondrogenesis"

The following table shows the presence or absence of the 14 vitamins or minerals (highlighted in yellow) which were determined to be absent in DMEM.

**Supplemental Table 1 – Common media compositions**

| **Medium** | **DMEM** | **RPMI-1640** | **Ham's F12** | **IMDM** |
|---|---|---|---|---|
| **Catalog number** | D5796 | R8758 | 51651C | I2762 |
| **Component (g/L)** | | | | |
| **Inorganic Salts** | | | | |
| Calcium Chloride | 0.2 | | 0.03322 | 0.219 |
| Calcium nitrate $Ca(NO_3)_2 \cdot 4H_2O$ | | 0.1 | | |
| Chromium | | | | |
| Cobalt (*present due to Vitamin B12) | | * | * | * |
| Copper Sulphate $CuSO_4 \cdot 9H_2O$ | | | 0.0000025 | |
| Ferric Nitrate • $9H_2O$ | 0.0001 | | 0.000834 | |
| Iodine | | | | |
| Manganese | | | | |
| Magnesium Sulfate (anhydrous) | 0.09767 | 0.04884 | | 0.09767 |
| Magnesium Chloride $MgCl_2$ | | | 0.05722 | |
| Molybdenum | | | | |
| Potassium Chloride | 0.4 | 0.4 | 0.224 | 0.33 |
| Potassium Nitrate $KNO_3$ | | | | 0.000076 |
| Sodium Bicarbonate $NaHCO_3$ | 3.7 | 2 | 1.176 | 3.024 |
| Sodium Chloride | 6.4 | 6 | 7.599 | 4.505 |
| Sodium Phosphate dibasic (anhydrous) $Na_2HPO_4$ | | 0.8 | 0.14204 | |
| Sodium Phosphate Monobasic (anhydrous) $NaH_2PO_4$ | 0.109 | | | 0.109 |
| Sodium Selenite $Na_2SeO_3$ | | | | 0.000017 |
| Zinc $ZnSO_4 \cdot 7H_2O$ | | | 0.000863 | |
| **Amino Acids** | | | | |
| L-Alanine | | | 0.009 | 0.025 |
| L-Arginine • HCl | 0.084 | 0.2 | 0.211 | 0.084 |
| L-Asparagine (Anhydrous) | | 0.05 | | 0.0284 |
| L-Asparagine • $H_2O$ | | | 0.01501 | |
| L-Aspartic Acid | | 0.02 | 0.0133 | 0.03 |
| L-Cystine • 2HCl | 0.0626 | 0.0652 | | 0.09124 |
| L-Cystine • HCl • $H_2O$ | | | 0.035 | |
| L-Glutamic Acid | | 0.02 | 0.0147 | 0.075 |
| L-Glutamine | 0.584 | 0.3 | 0.146 | |
| Glycine | 0.03 | 0.01 | 0.00751 | 0.03 |
| L-Histidine (free base) | | 0.015 | | |
| L-Histidine • HCl • $H_2O$ | 0.042 | | 0.02096 | 0.042 |
| L-Isoleucine | 0.105 | 0.05 | 0.00394 | 0.105 |
| L-Leucine | 0.105 | 0.05 | 0.0131 | 0.105 |
| L-Lysine • HCl | 0.146 | 0.04 | 0.0365 | 0.146 |
| L-Methionine | 0.03 | 0.015 | 0.00448 | 0.03 |
| L-Phenylalanine | 0.066 | 0.015 | 0.00496 | 0.066 |

| | | | | |
|---|---|---|---|---|
| L-Proline | | 0.02 | 0.0345 | 0.04 |
| L-Serine | 0.042 | 0.03 | 0.0105 | 0.042 |
| L-Threonine | 0.095 | 0.02 | 0.0119 | 0.095 |
| L-Tryptophan | 0.016 | 0.005 | 0.0024 | 0.016 |
| L-Tyrosine • 2Na •$2H_2O$ | 0.10379 | 0.02883 | 0.00778 | 0.10379 |
| L-Valine | 0.094 | 0.02 | 0.0117 | 0.094 |
| **Vitamins** | | | | |
| Choline Chloride | 0.004 | 0.003 | 0.01396 | 0.004 |
| *myo*-Inositol | 0.0072 | 0.035 | 0.018 | 0.0072 |
| Vitamin A | | | | |
| Vitamin B1 Thiamine • HCl | 0.004 | 0.001 | 0.000034 | 0.004 |
| Vitamin B2 Riboflavin | 0.0004 | 0.0002 | 0.000038 | 0.0004 |
| Vitamin B3 Niacinamide | 0.004 | 0.001 | 0.0000367 | 0.004 |
| Vitamin B5 D-Pantothenic Acid (hemicalcium) | 0.004 | 0.00025 | 0.000238 | 0.004 |
| Vitmain B6 Pyridoxal • HCl | — | | | 0.004 |
| Vitamin B6 Pyridoxine • HCl | 0.00404 | 0.001 | 0.000062 | |
| Vitamin B7 D-Biotin | | 0.0002 | 0.0000073 | 0.000013 |
| Vitamin B9 Folic Acid | 0.004 | 0.001 | 0.0013 | 0.004 |
| Vitamin B12 | | 0.000005 | 0.00136 | 0.000013 |
| Vitamin D | | | | |
| Vitamin E | | | | |
| Vitamin K | | | | |
| **Other** | | | | |
| p-Amino Benzoic Acid | | 0.001 | | |
| D-Glucose | 4.5 | 2 | 1.802 | 4.5 |
| Glutathione (reduced) | | 0.001 | | |
| HEPES | | | | 5.958 |
| Hypoxanthine sodium salt | | | 0.00477 | |
| Linoleic Acid | | | 0.000084 | |
| α-linolenic acid | | | | |
| Phenol Red • Na | 0.0159 | 0.0053 | 0.0013 | 0.016 |
| Pyruvic Acid • Na | | | 0.11 | 0.11 |
| Putrescine •2H2O | | | 0.000161 | |
| Thioctic Acid | | | 0.00021 | |
| Thymidine | | | 0.00073 | |
| **Add** | | | | |
| L-Glutamine | | | | 0.584 |

**Supplemental Data 1**

The following graphs all show type II collagen expression, assessed by luminescence, at 4 time points following continuous stimulation with vitamins or minerals over a 5 log dose response ($n \geq 3 \pm$ S.D., * > control, basal medium, * < control, basal medium; 2-way ANOVA with Sidak's multiple comparison test, Alpha 0.05). In all comparisons both day and dose were significant sources of variation with significant interaction between them ($p<0.0001$).

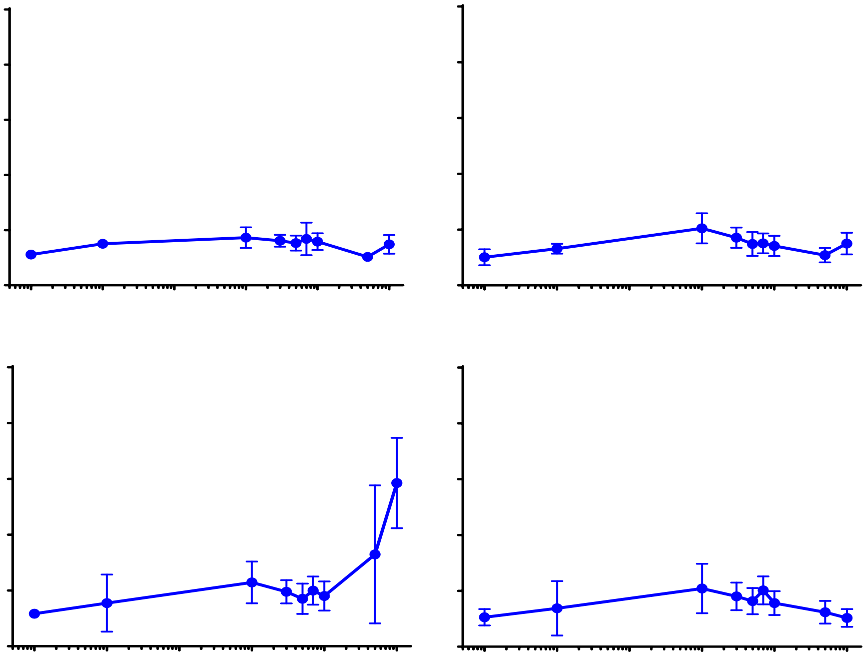

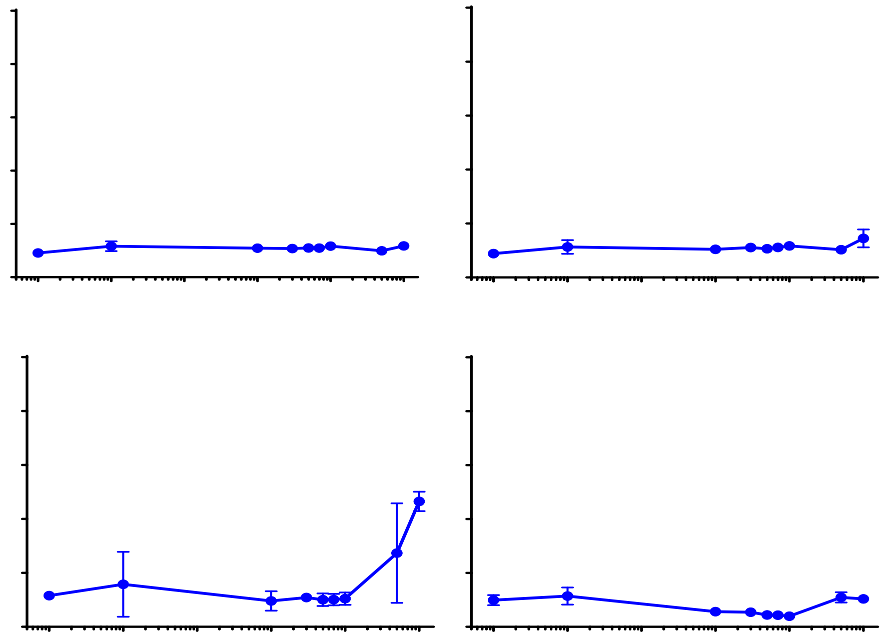

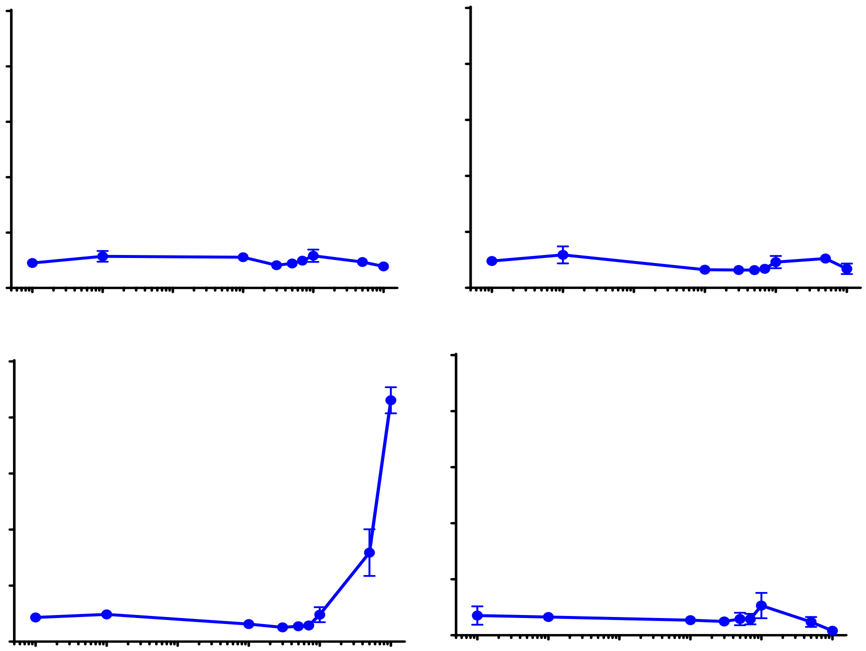

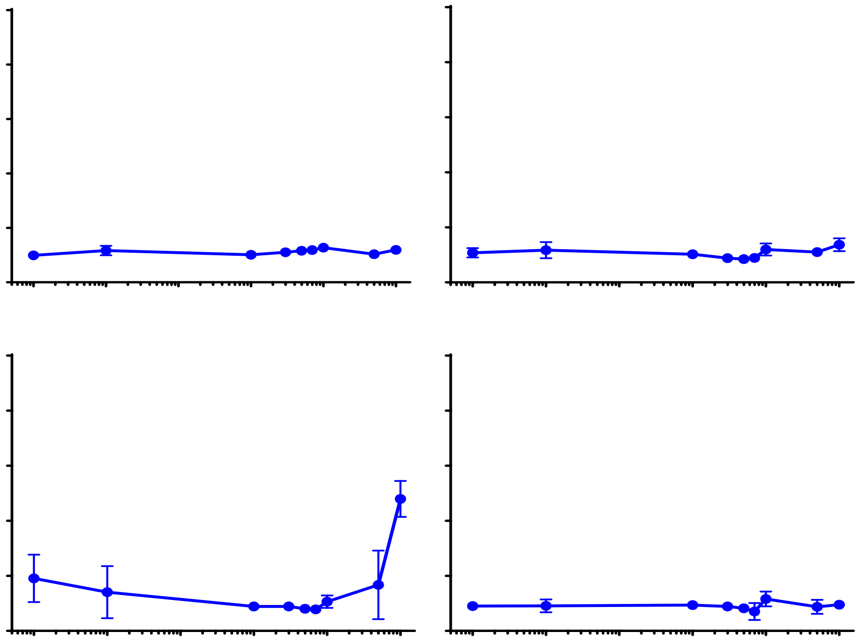

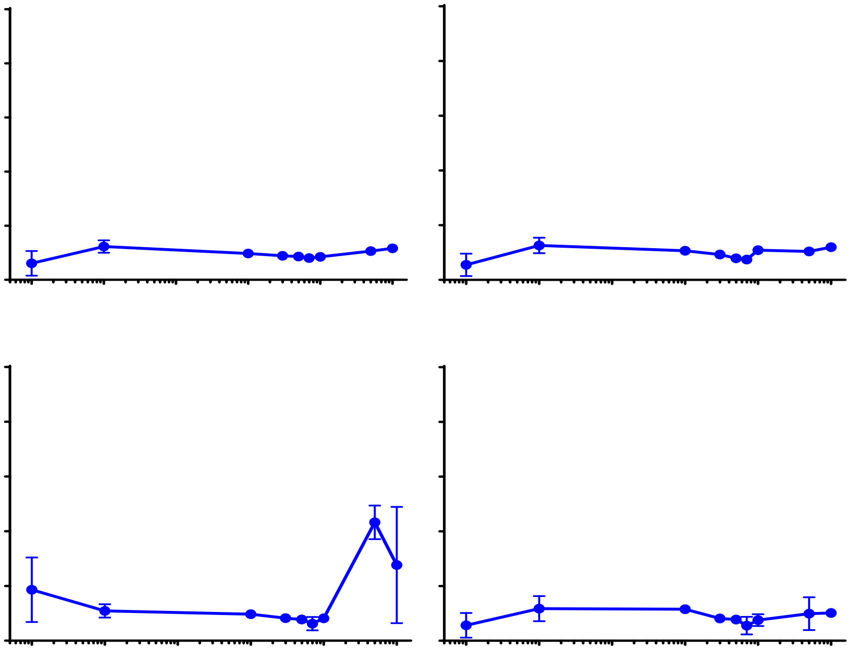

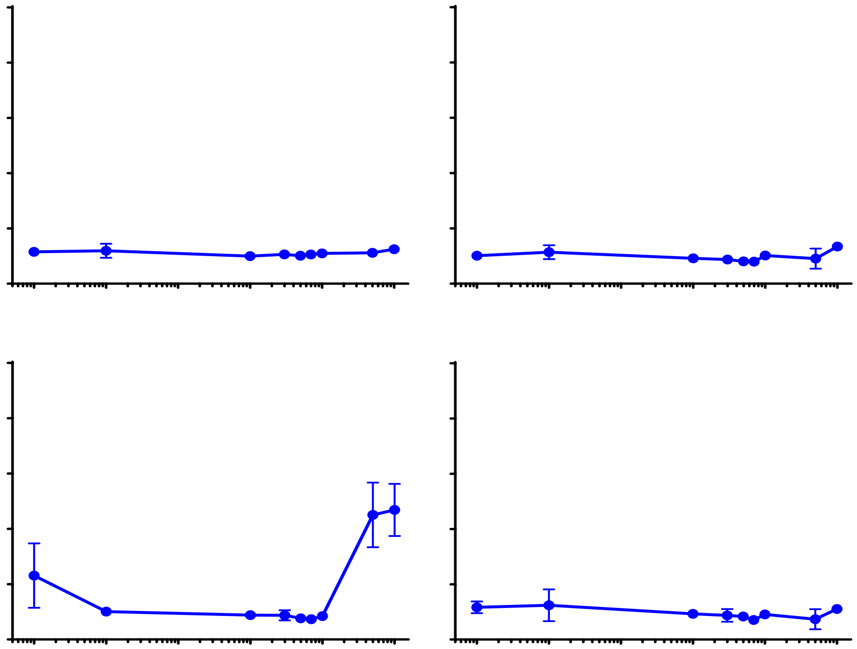

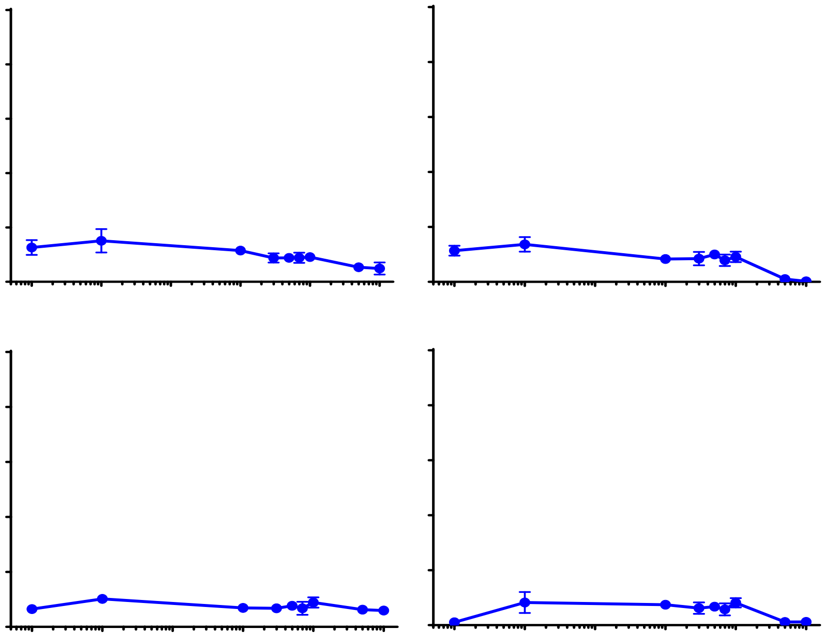

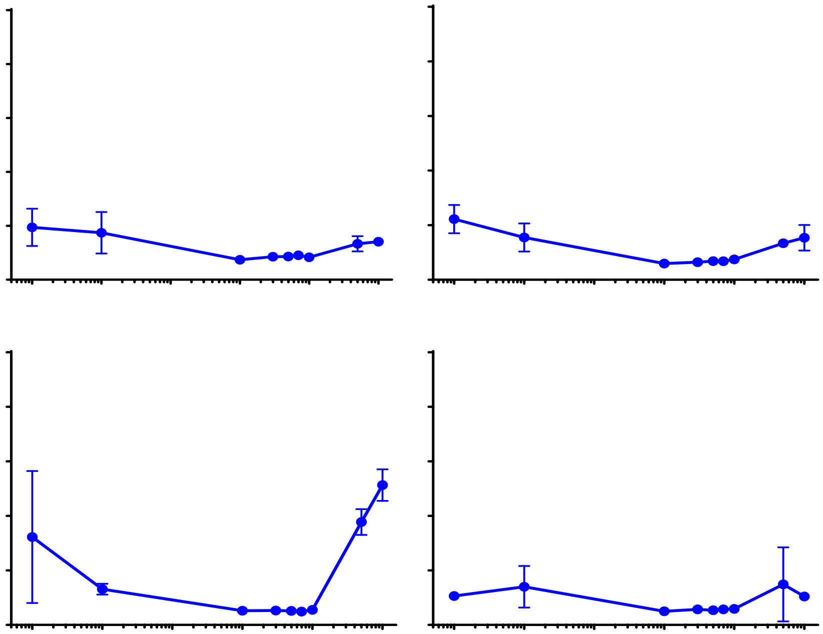

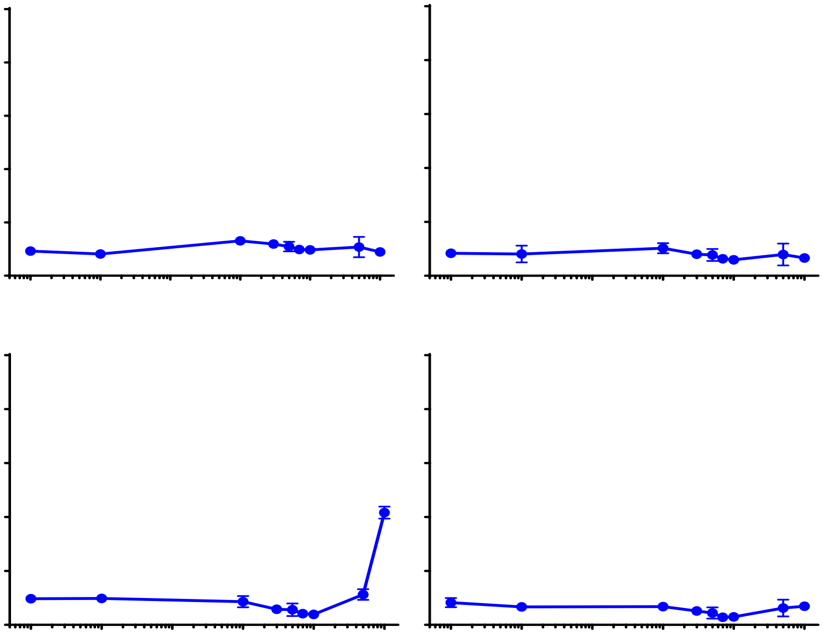

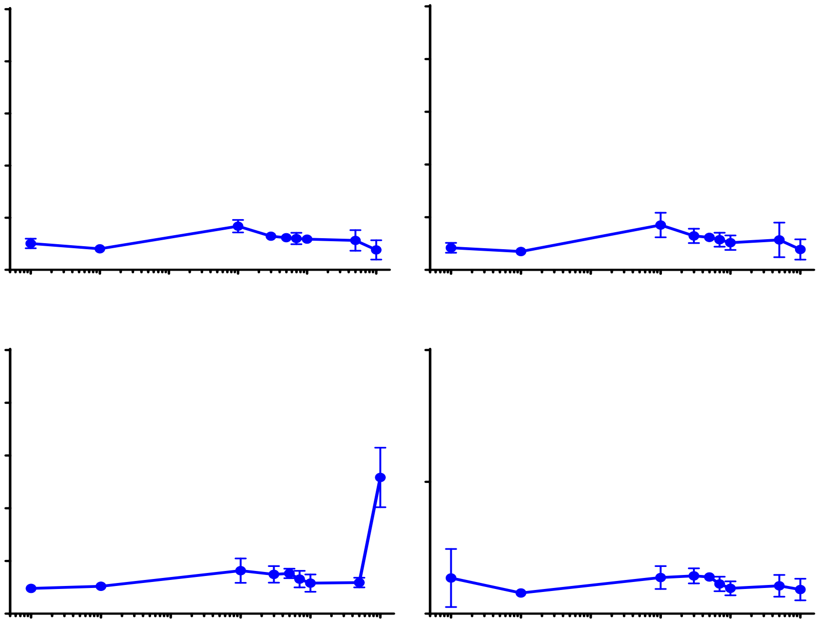

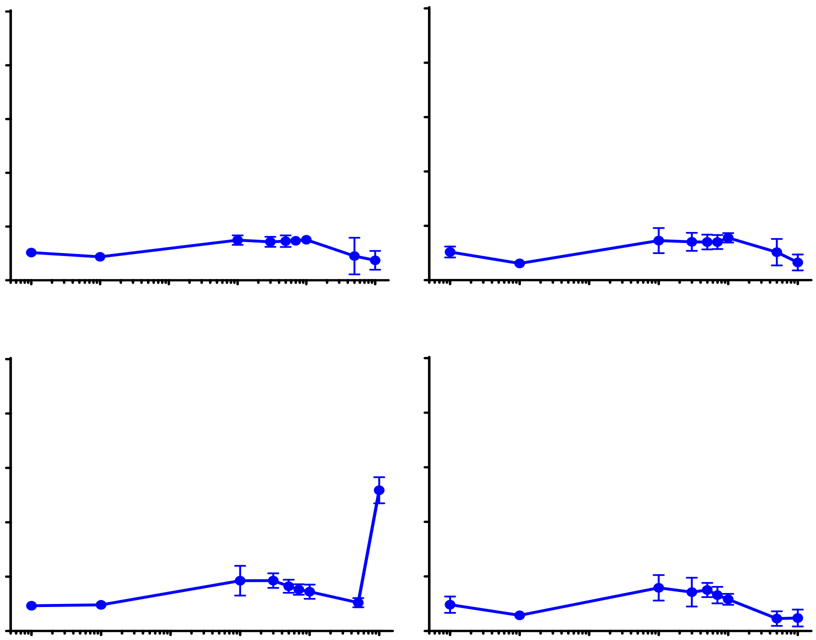

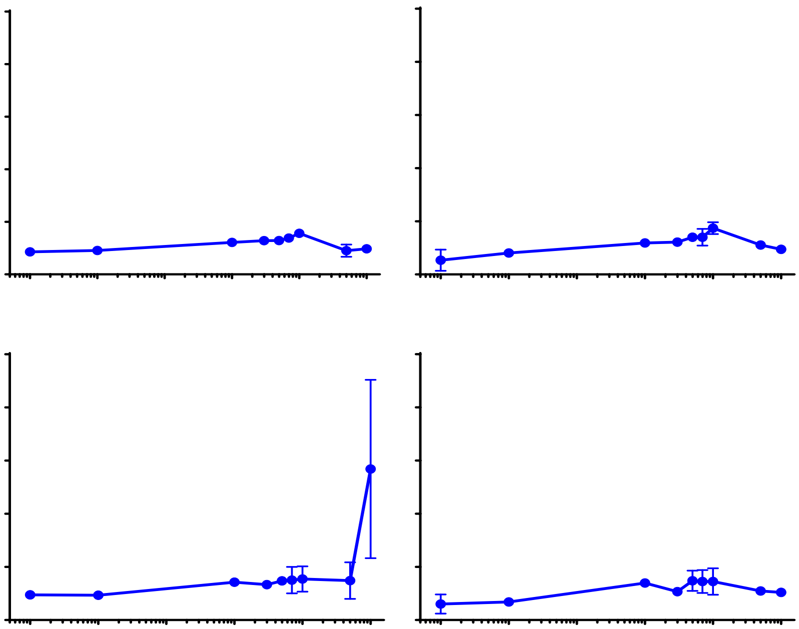

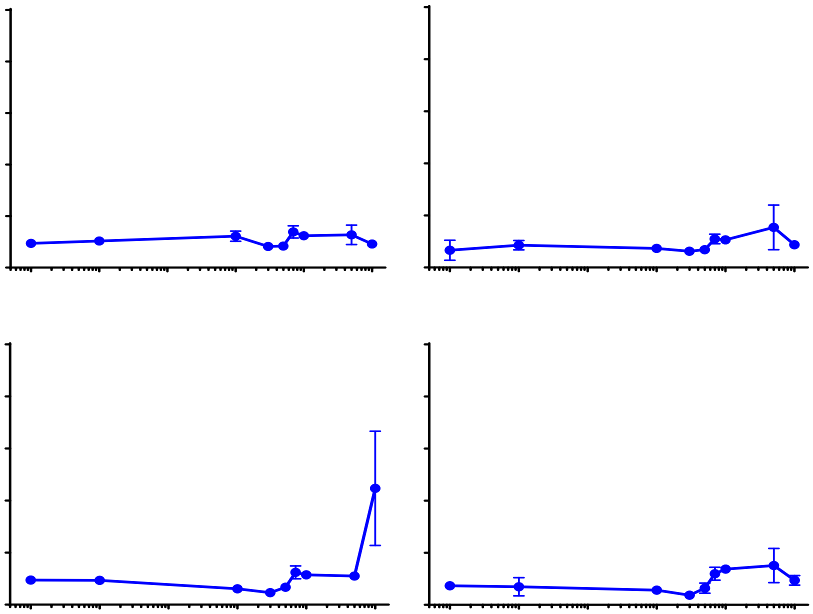

**Supplemental Data 2**

The following graphs show the results of supplementation of TGFβ1 containing media with vitamins and minerals alone or in combination. Media were normalized against basal control medium with no TGFβ1 and compared with that and the TGFβ1 supplemented medium by t-test with false discovery rate set at 1%. TGFβ1 supplemented medium is represented by a red dashed line. Conditions that were statistically greater than TGFβ1 supplemented medium are represented by a ▲ and an *; conditions that are greater than basal but not TGFβ1 are represented by ●; conditions that are greater than basal but lower than TGFβ1 are represented by ▼ and an L; conditions that were not significantly different to basal are represented by ■ (these are also significantly lower than TGFβ1 after 0.4 week); there were no conditions that were significantly lower than basal medium alone.

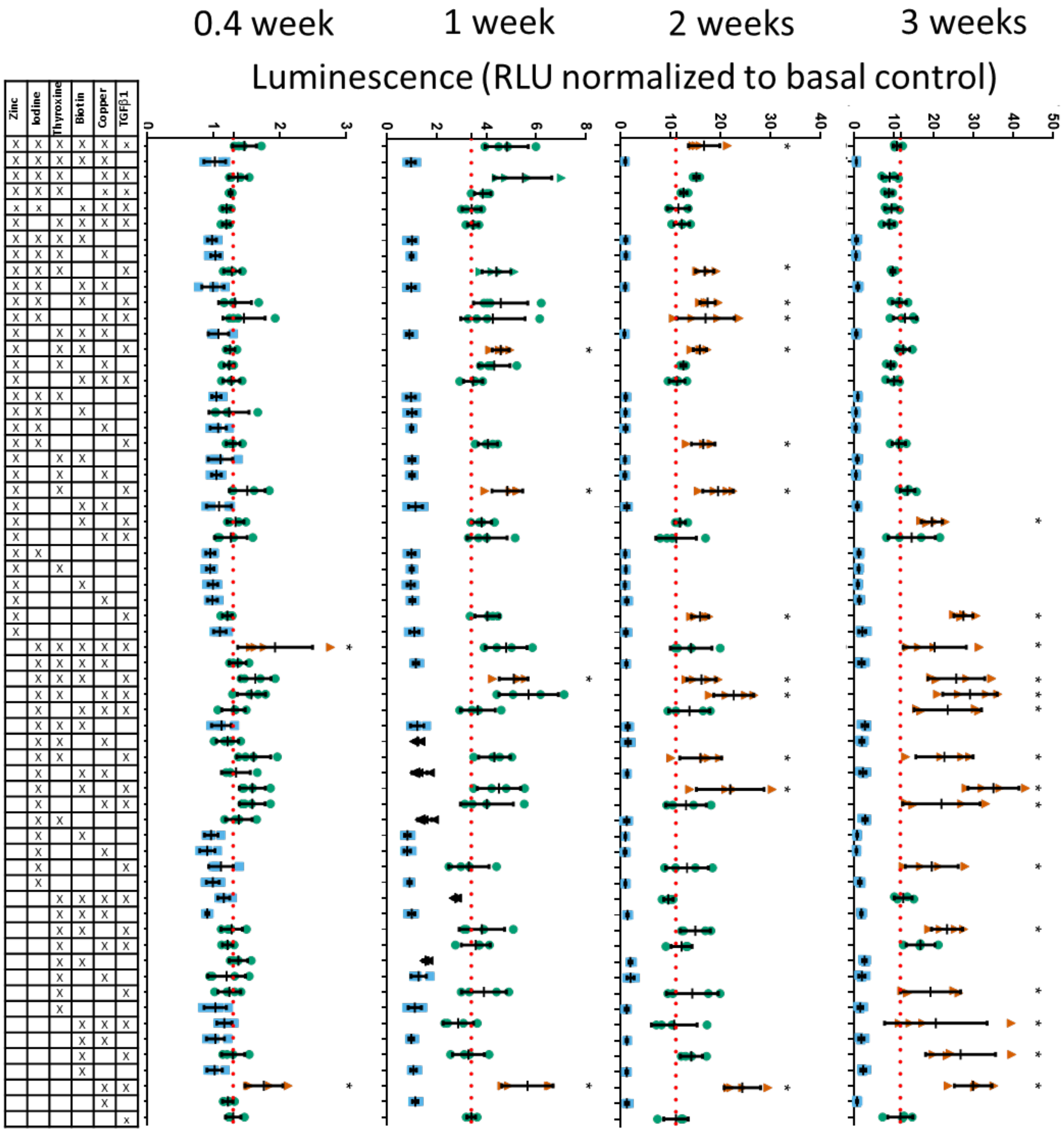

**Supplemental Data 3**

The following graphs show the results of addition of vitamins B12, E, and chromium to previously defined supplemented media over a three-week period. Media were normalized against basal control medium with no TGFβ1 and compared to each supplemented media by two-way ANOVA with Sidak's multiple comparison test, Alpha 0.05. Each supplemented media average is represented by a red dashed line. Conditions that were statistically greater than their respective supplemented medium are represented by a ▲ and an *; conditions that were similar to their respective supplemented medium are represented by a ●; conditions that were lower than their respective supplemented medium are represented by ▼. There was significant interaction between the supplemented media and the additional factors, both of which were also significant sources of variation.

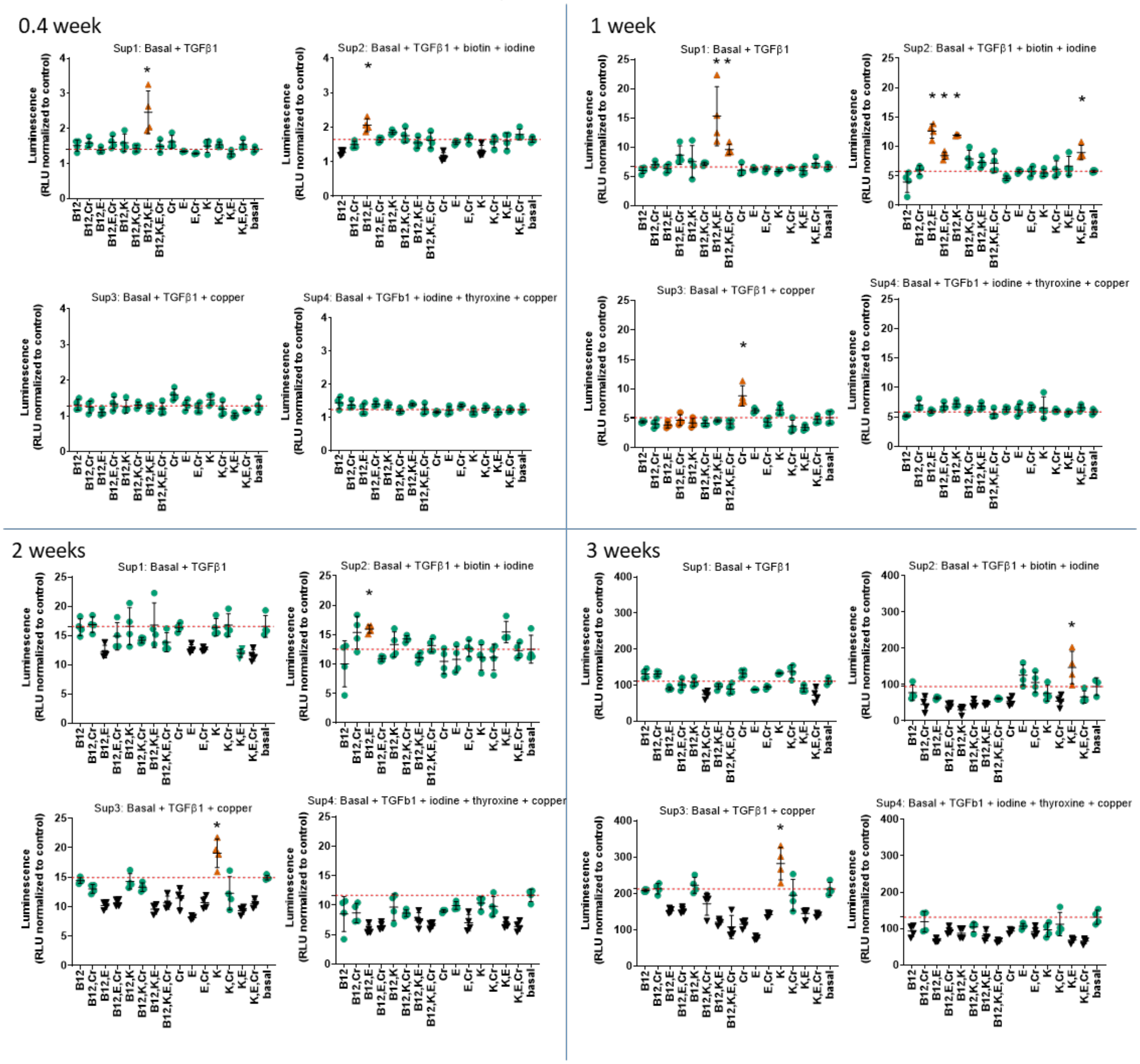

**Supplemental Data 4**

The following graphs all show type II collagen expression, assessed by luminescence, at 4 time points following continuous stimulation with vitamins or minerals in the presence of TGFβ1 (1ng/ml) over a 5 log dose response. Conditions that were statistically greater than TGFβ1 supplemented medium are represented by a ▲ and an *; conditions that were similar to TGFβ1 supplemented medium are represented by a ●; conditions that were lower than TGFβ1 supplemented medium are represented by ▼; 2-way ANOVA with Sidak's multiple comparison test, Alpha 0.05. A red dashed line indicates the mean TGFβ1 supplemented medium response. In most comparisons both time and dose were significant sources of variation with significant interaction between them, where this was not the case details are noted below the set of graphs.

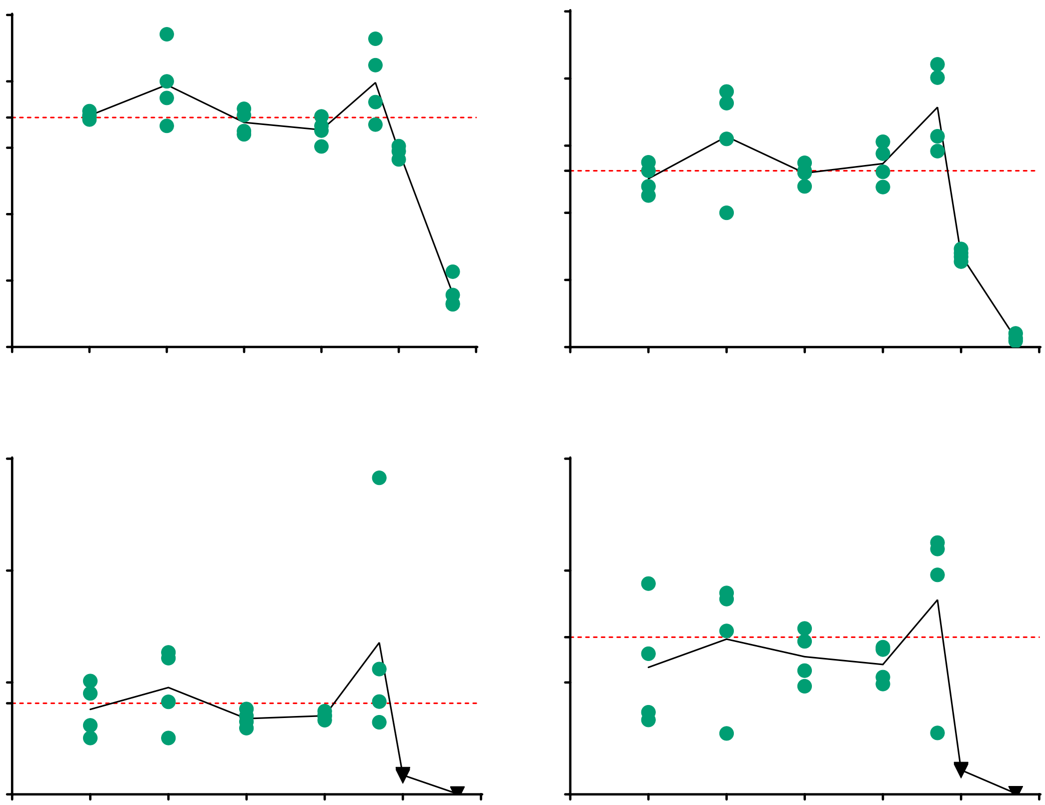

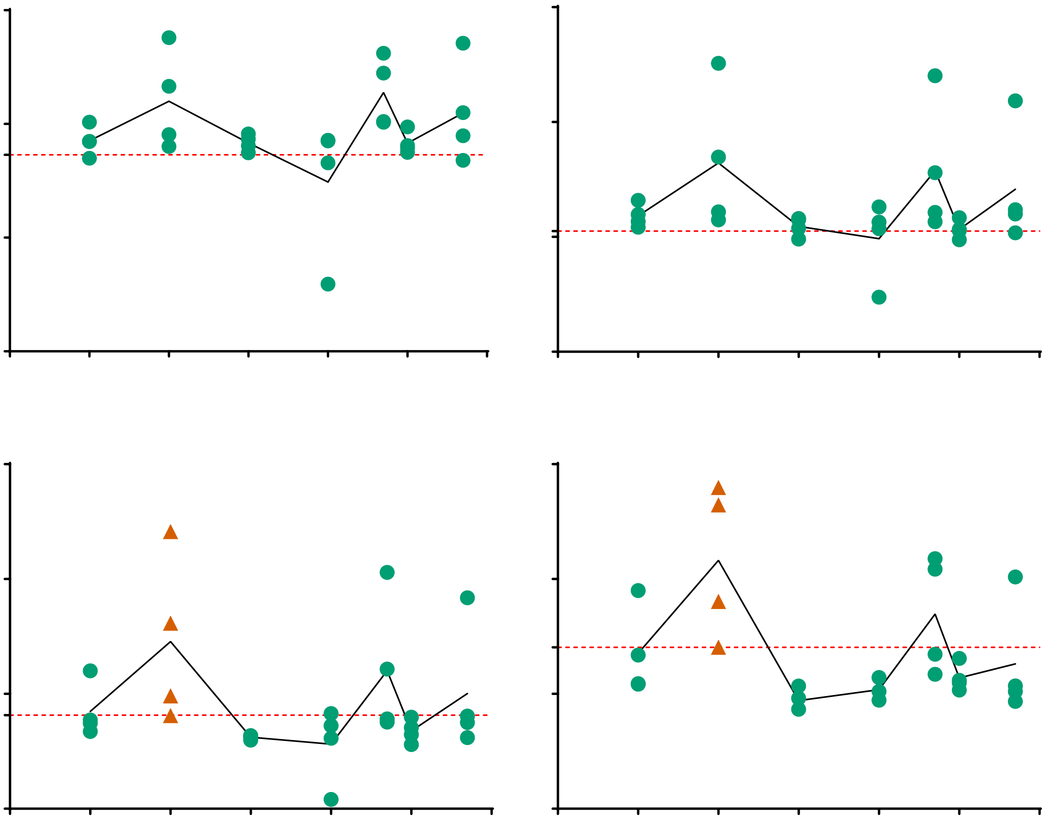

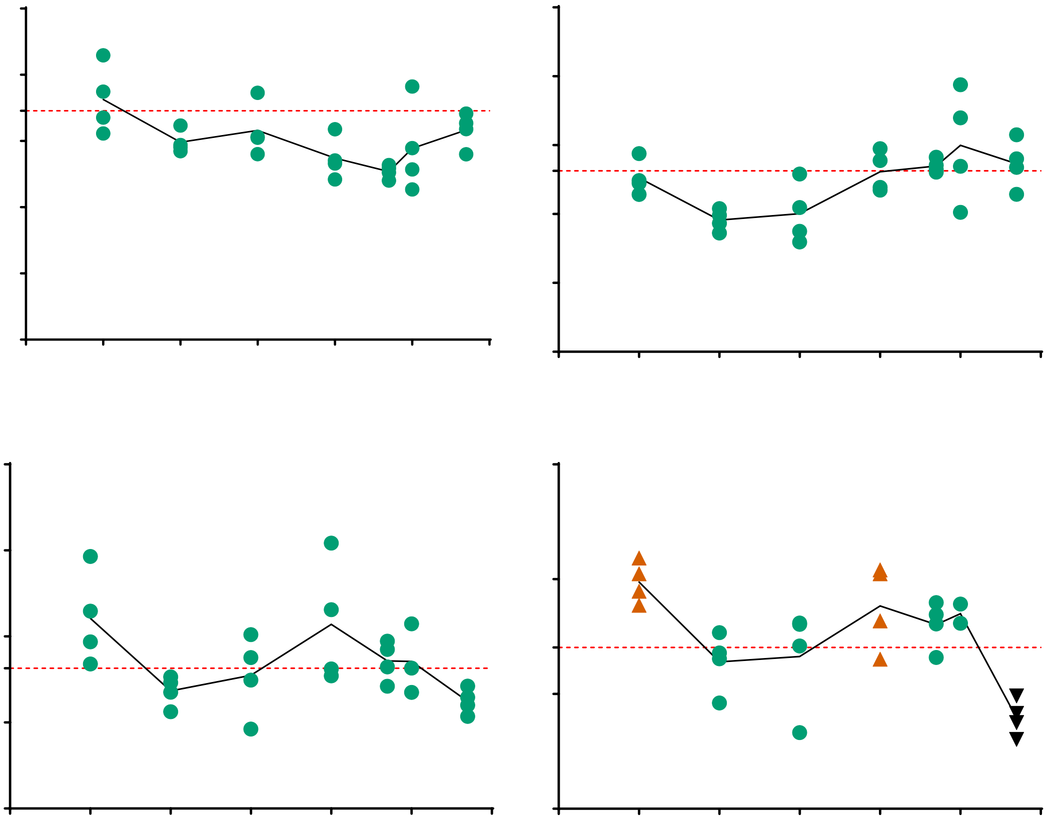

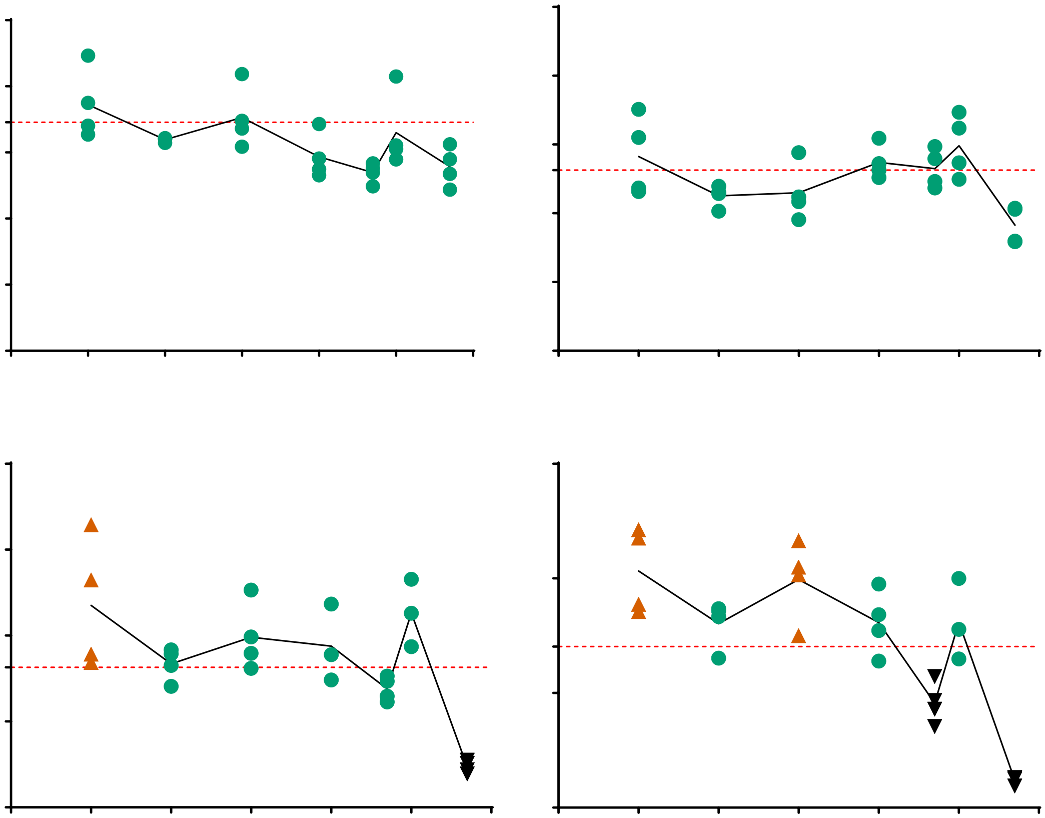

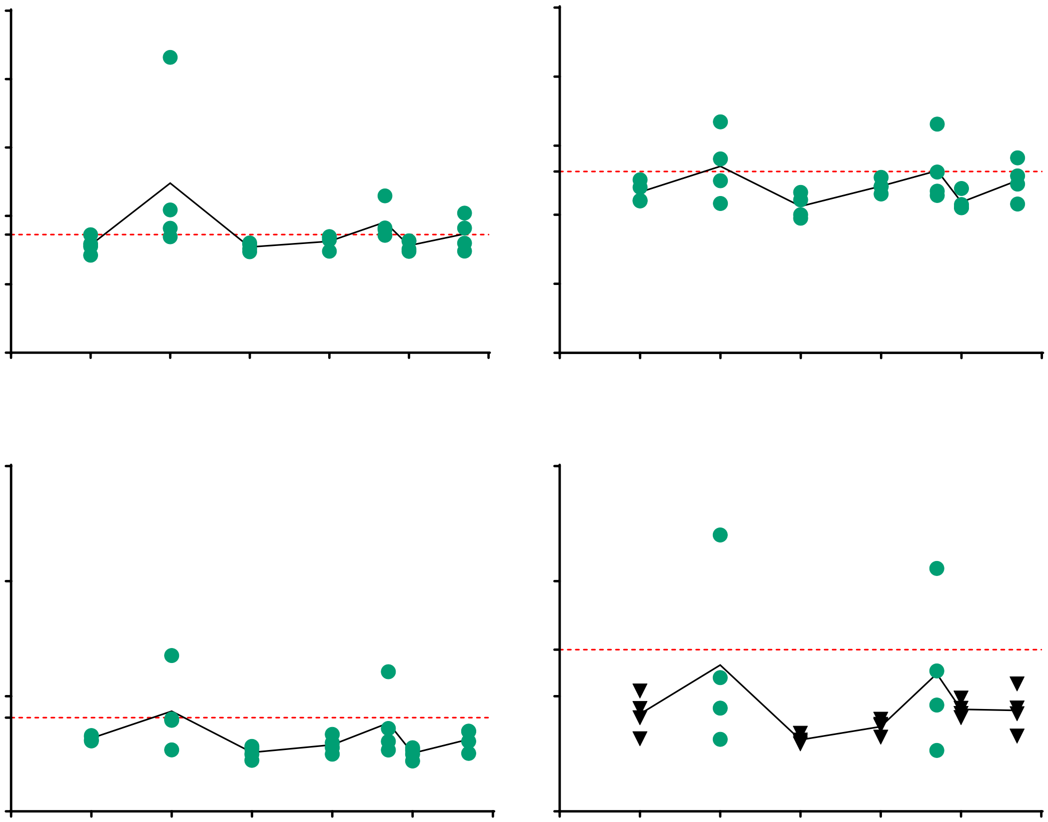

There was no significant interaction found between time and dose, both time ($p < 0.0001$) and dose ($p = 0.0007$) were significant sources of variation.

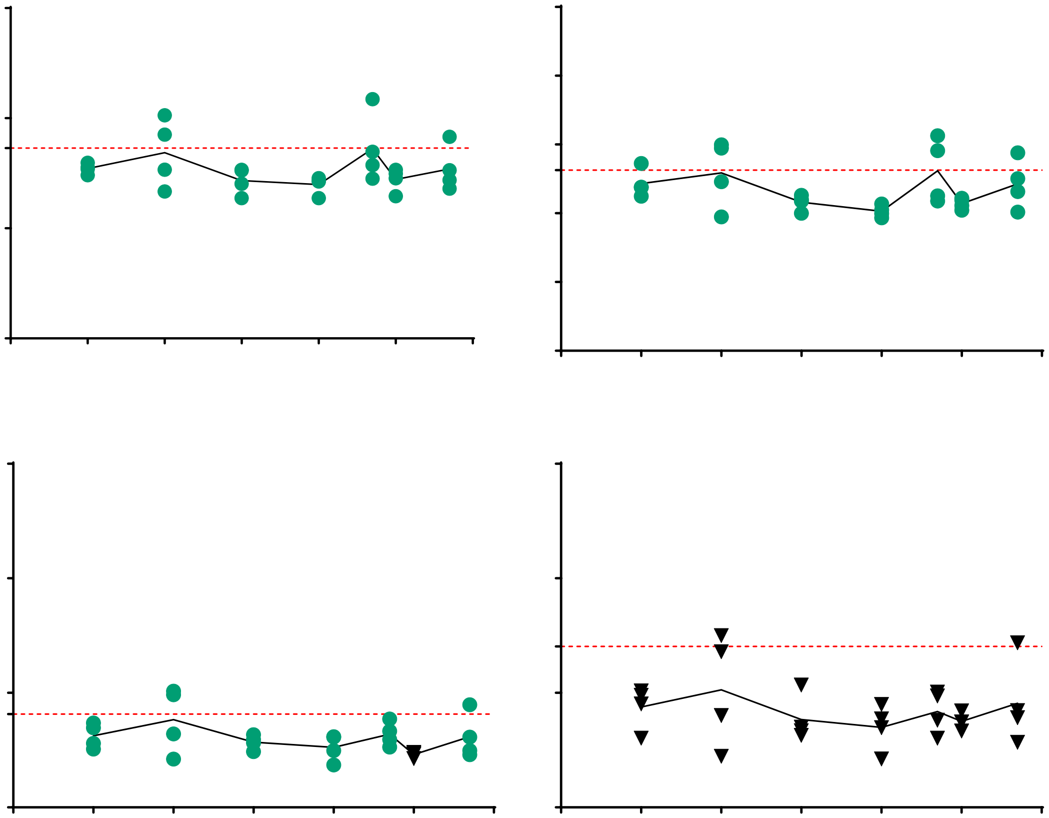

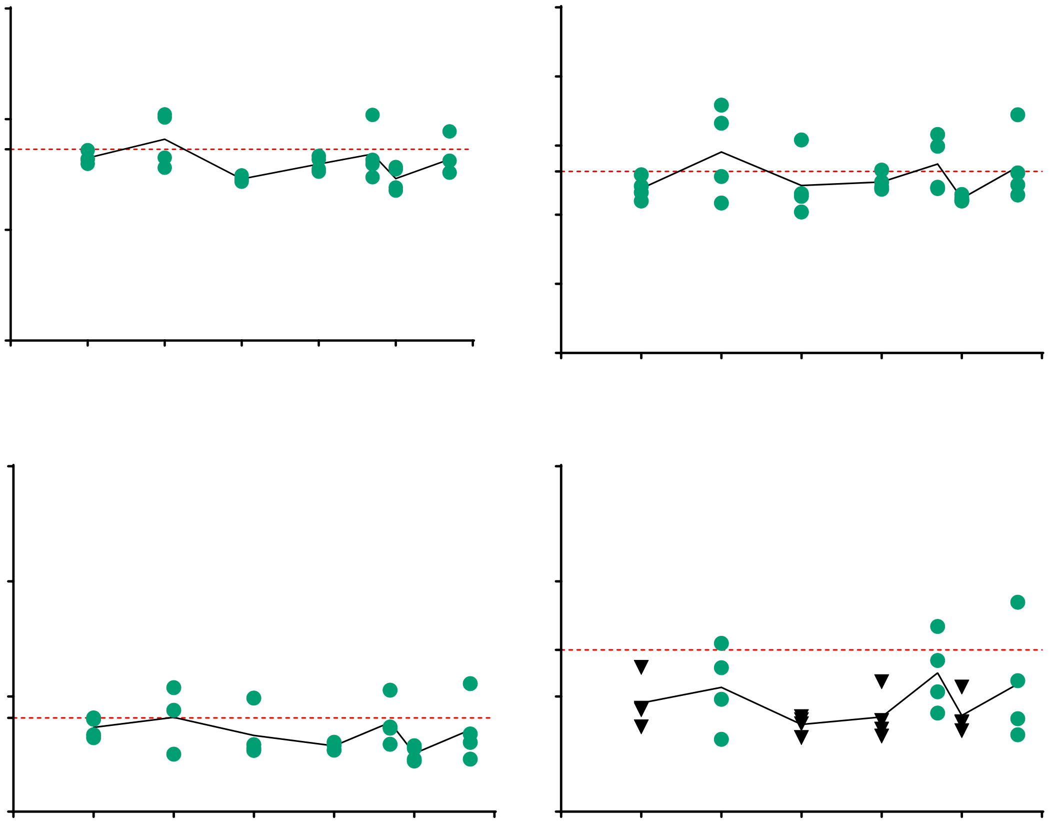

There was no significant interaction found between time and dose; both time ($p < 0.0001$) and dose ($p = 0.0024$) were significant sources of variation.

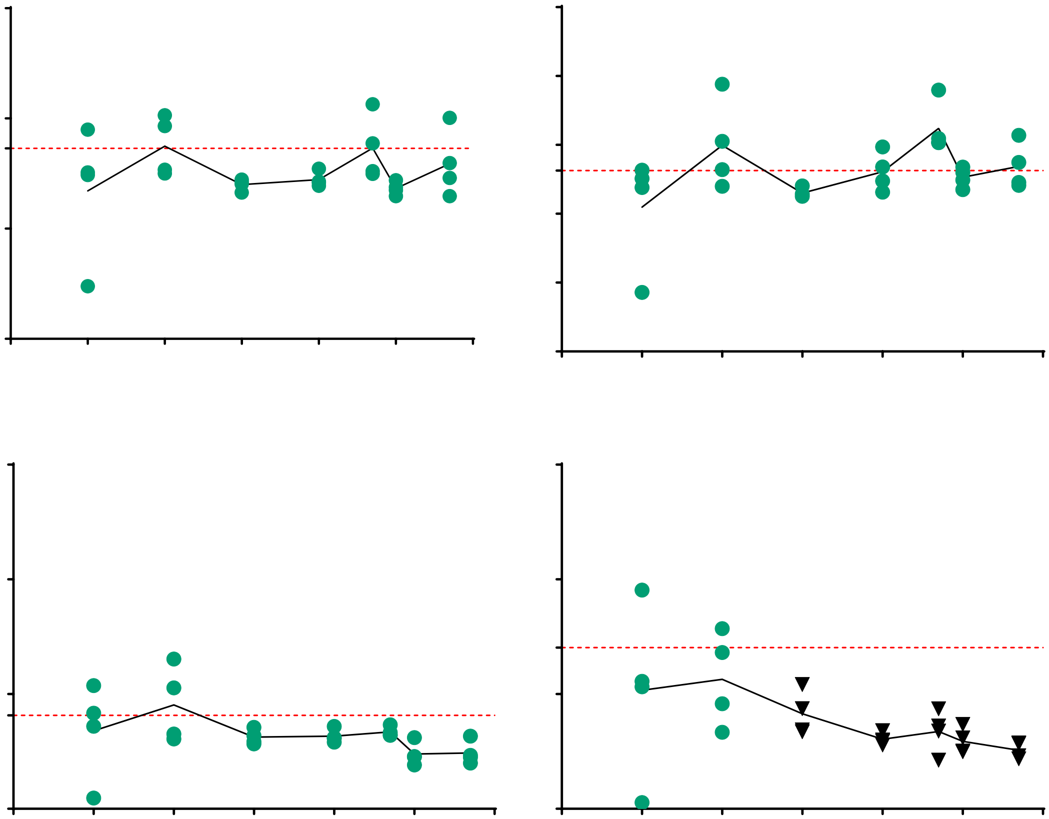

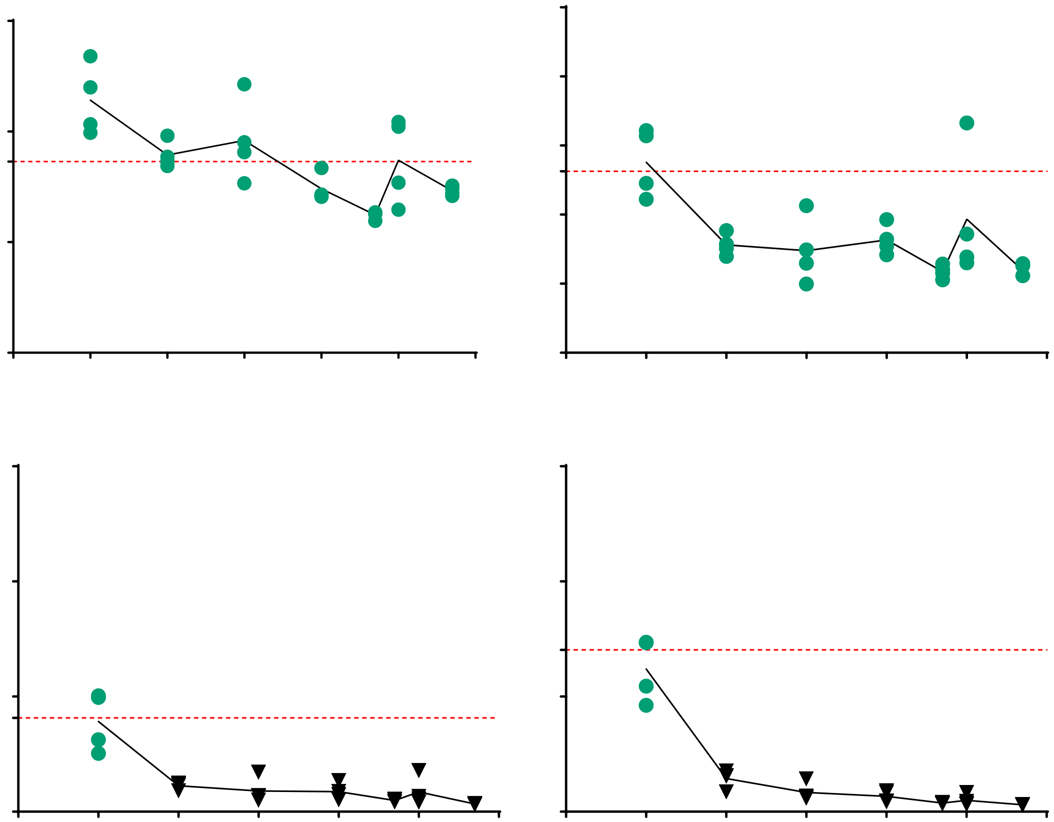

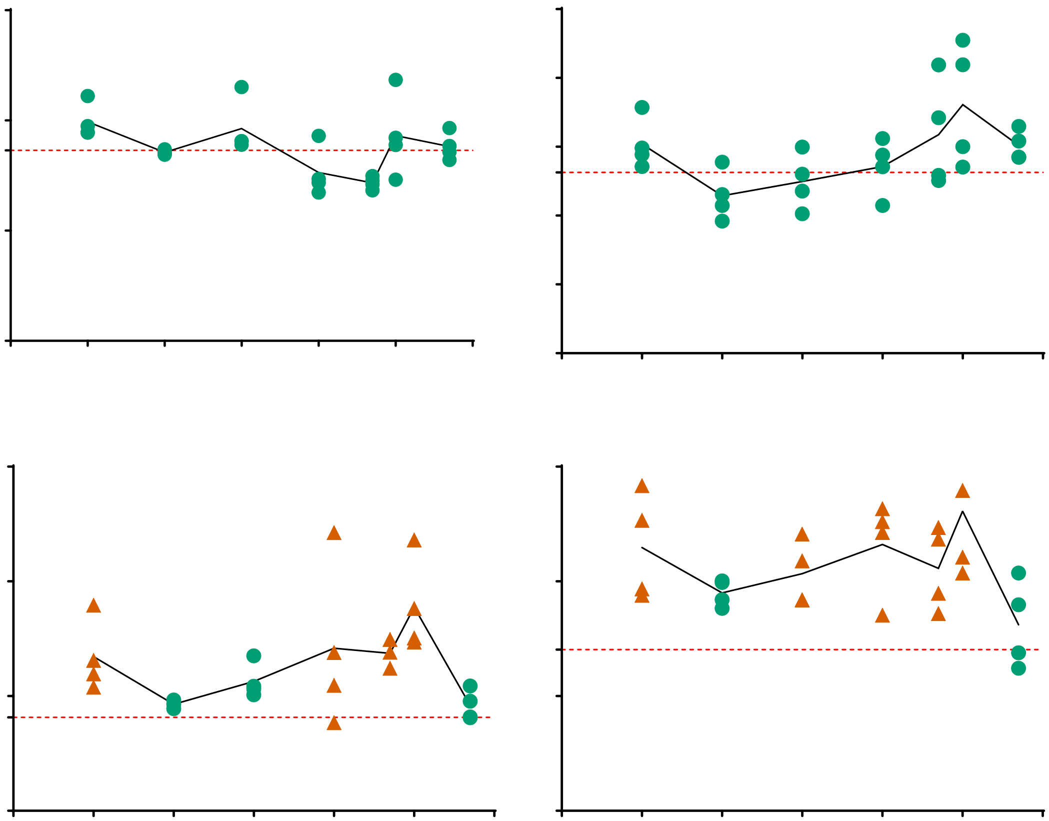

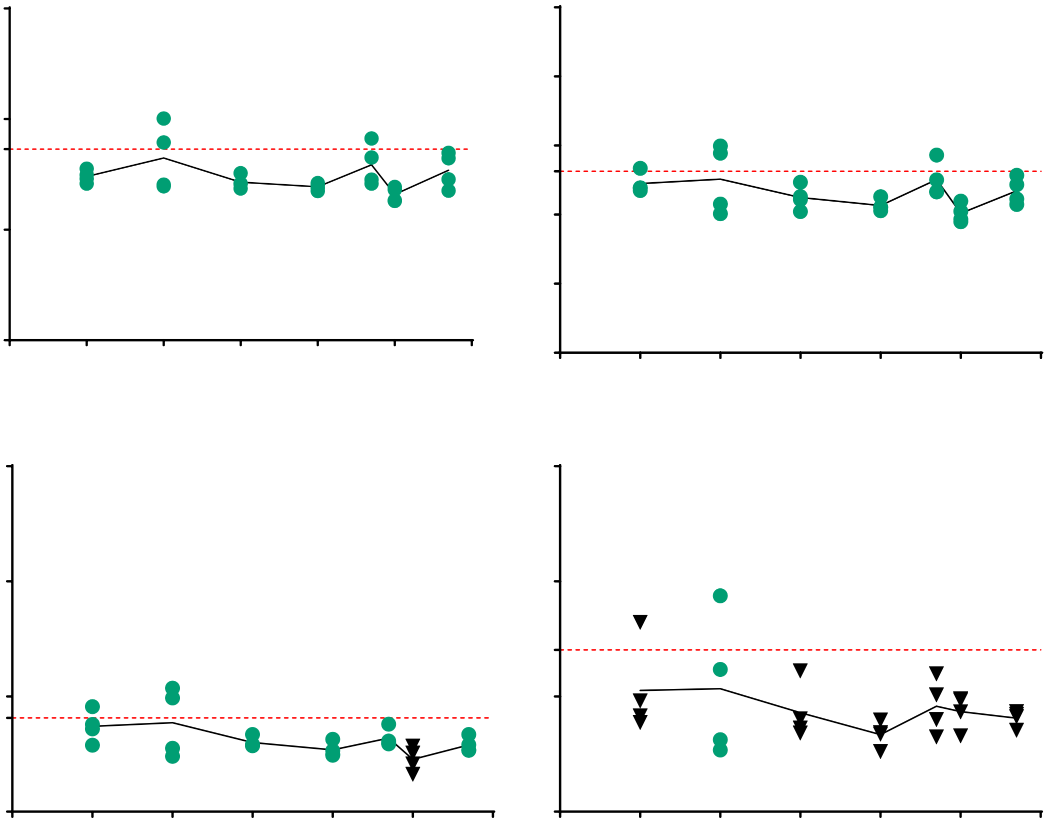

There was no significant interaction found between time and dose; both time ($p < 0.0001$) and dose ($p = 0.0005$) were significant sources of variation.

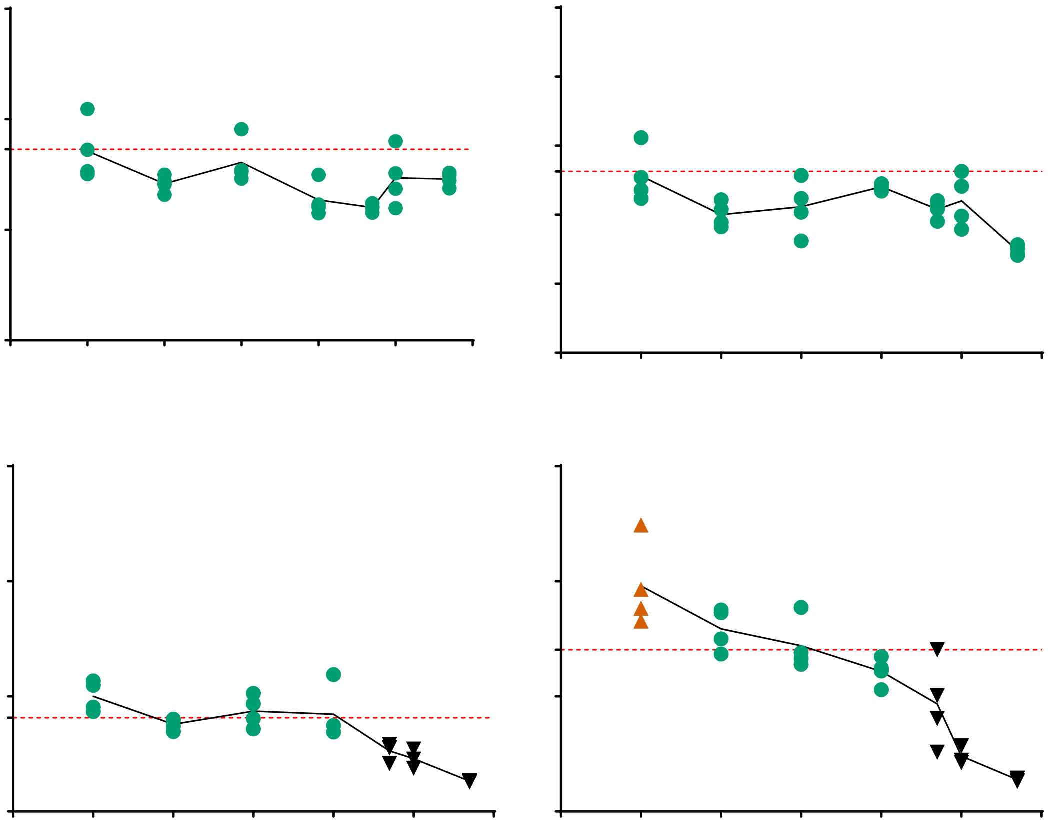

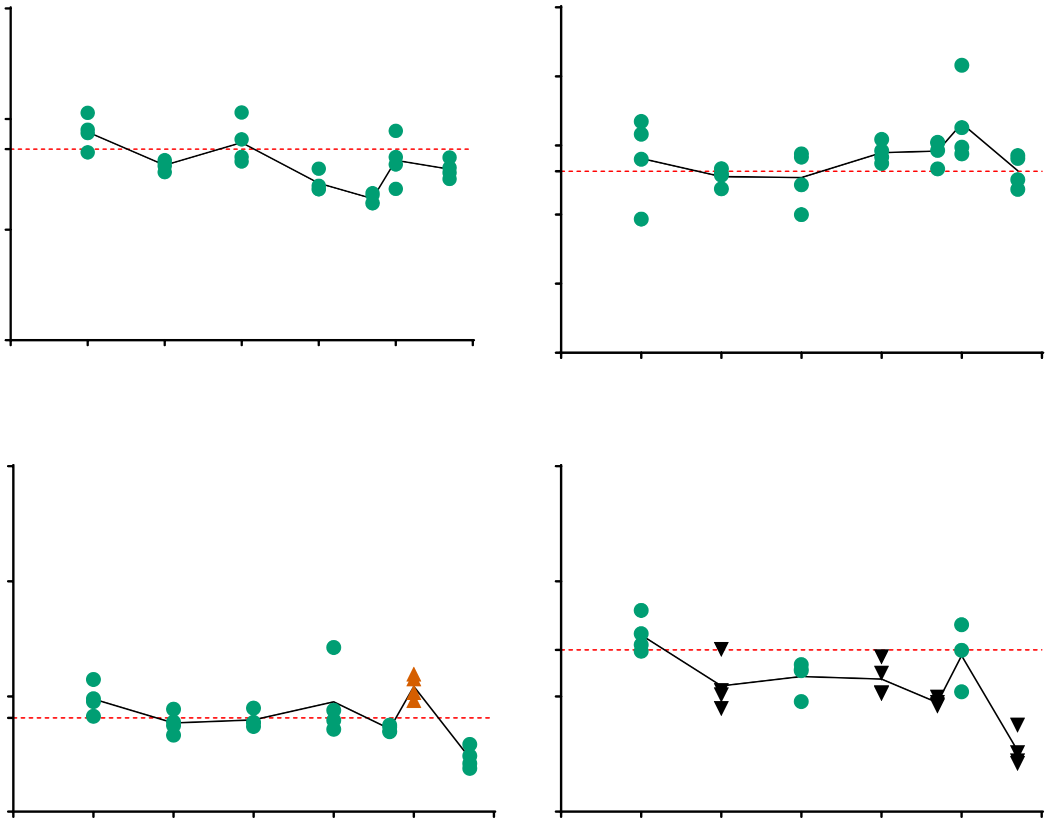

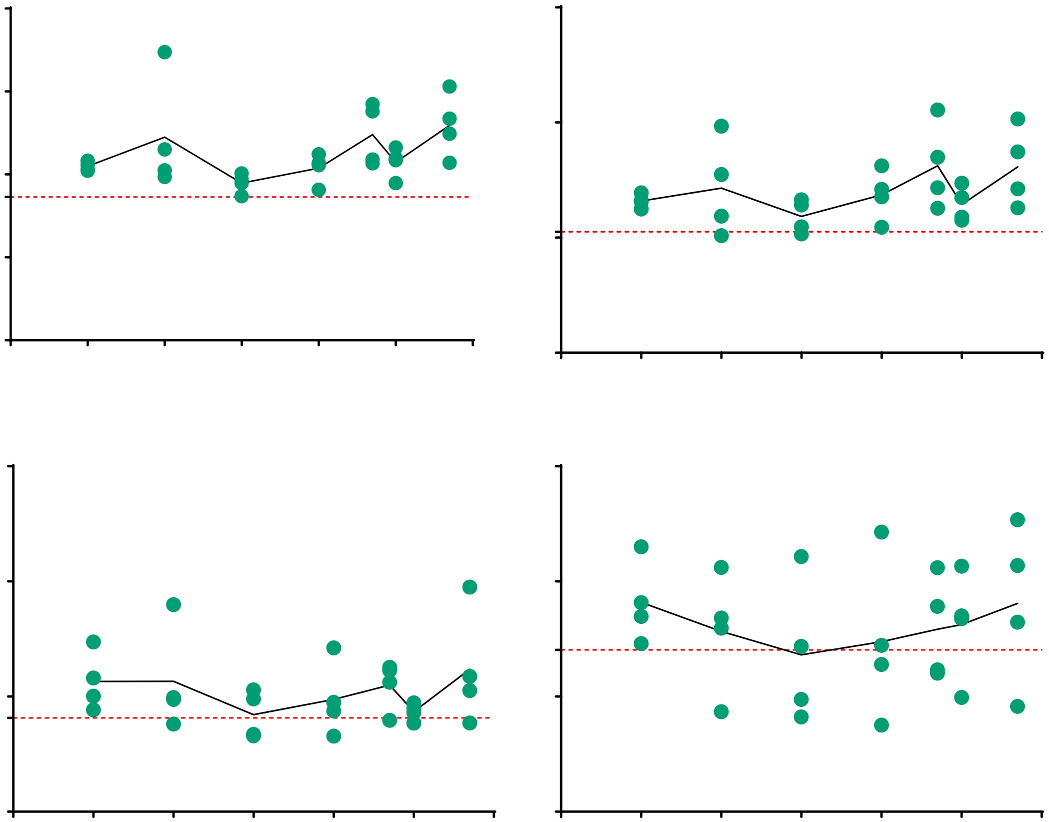

There was no significant interaction found between time and dose; only time ($p < 0.0001$) was a significant source of variation.

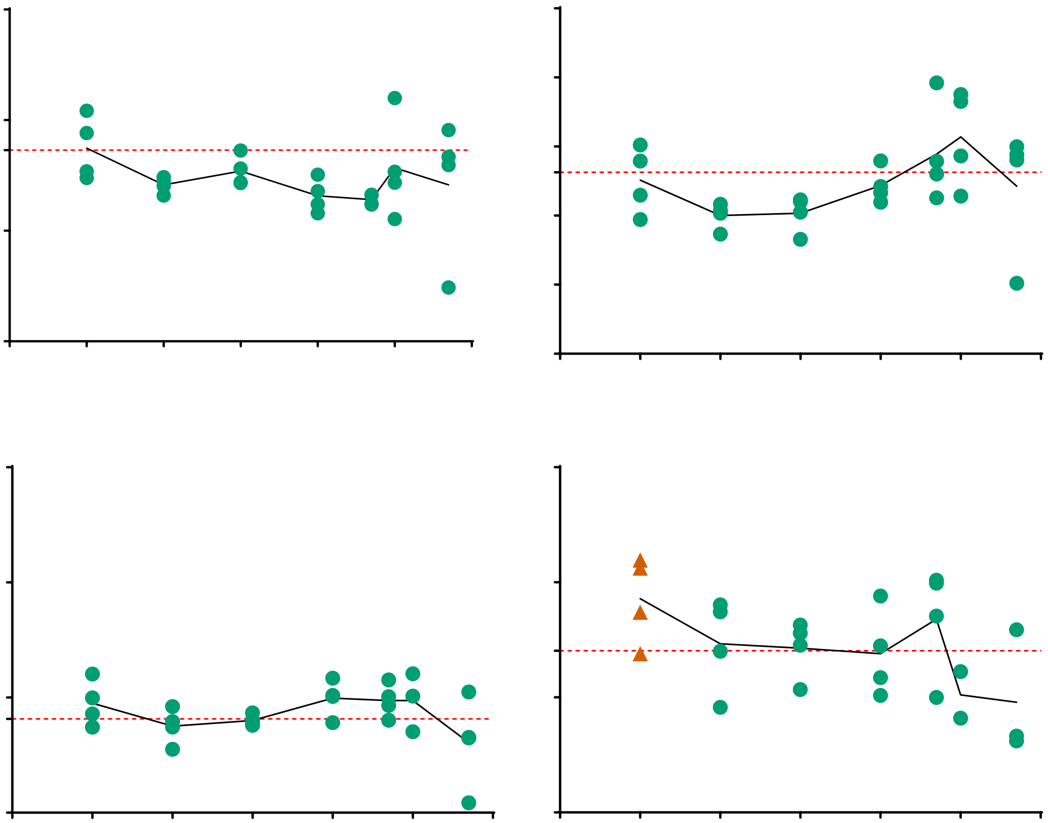

**Supplemental Data 5**

The following graphs all show metabolic activity, assessed by resazurin fluorescence, at day 21 following continuous stimulation with vitamins or minerals in the presence of TGFβ1 (1ng/ml) over a 5 log dose response. Conditions that were statistically greater than TGFβ1 supplemented medium are represented by a ▲ and an *; conditions that were similar to TGFβ1 supplemented medium are represented by a ●; conditions that were lower than TGFβ1 supplemented medium are represented by ▼; 1-way ANOVA with Dunnett's multiple comparison test, Alpha 0.05. A red dashed line indicates the mean TGFβ1 supplemented medium response.

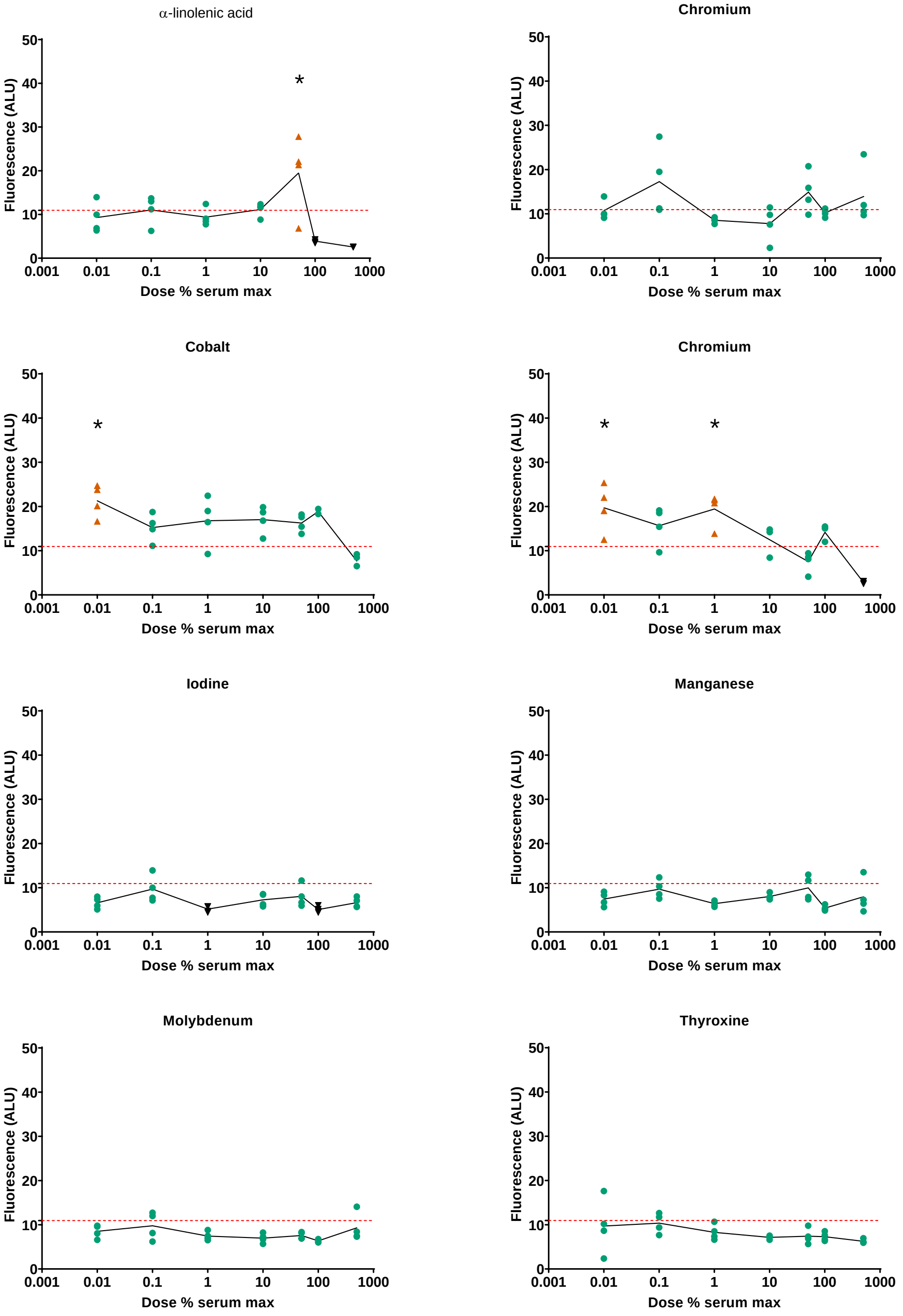

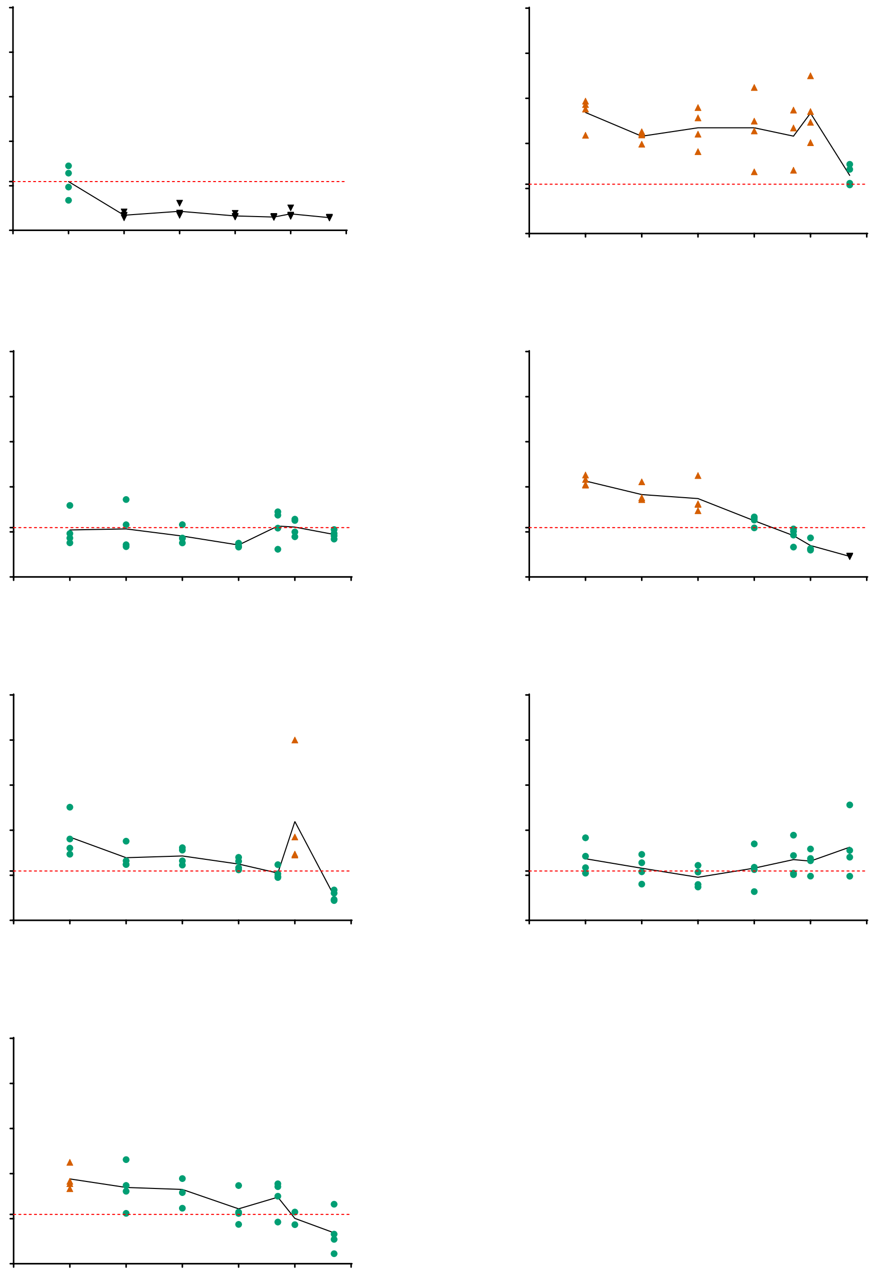

**Supplemental Data 6**

The following table shows the cumulative response (normalized to basal medium control) to vitamin and mineral supplementation over the 21-day duration of the experiment in the presence of TGF$\beta$1 (1ng/ml), the table is sorted in order of the highest net area.

| Factor (dose % serum max) | Net Area | Std. Error | 95% Confidence Interval |
|---|---|---|---|
| Vitamin B12 (100) | 434.3 | 55.07 | 326.4 to 542.3 |
| Chromium (0.1) | 369.1 | 84.07 | 204.3 to 533.9 |
| Vitamin B12 (10) | 355.2 | 76.11 | 206.0 to 504.4 |
| Vitamin B12 (0.01) | 348.2 | 42.19 | 265.5 to 430.9 |
| Vitamin B12 (50) | 341.9 | 26.09 | 290.8 to 393.0 |
| $\alpha$-linolenic acid (50) | 323 | 111.3 | 104.9 to 541.2 |
| Vitamin K (500) | 317.3 | 66.09 | 187.8 to 446.8 |
| Copper (0.01) | 308.1 | 44.43 | 221.0 to 395.2 |
| Chromium (50) | 303.8 | 69.33 | 167.9 to 439.7 |
| Vitamin B12 (1) | 297.7 | 22.98 | 252.6 to 342.7 |
| Vitamin K (0.01) | 293.1 | 33.25 | 227.9 to 358.2 |
| Cobalt (0.01) | 289.1 | 29.24 | 231.8 to 346.4 |
| Vitamin K (50) | 284.7 | 35.1 | 215.9 to 353.5 |
| Vitamin K (0.1) | 282.2 | 56.15 | 172.1 to 392.2 |
| Copper (100) | 276.5 | 29.24 | 219.2 to 333.8 |
| Vitamin D (0.01) | 273.9 | 26.17 | 222.6 to 325.2 |
| Cobalt (10) | 271.9 | 42.28 | 189.1 to 354.8 |
| Copper (1) | 271.2 | 29.04 | 214.2 to 328.1 |
| Zinc (0.01) | 260.2 | 30.43 | 200.6 to 319.8 |
| Vitamin E (100) | 258.6 | 21.22 | 217.0 to 300.2 |
| Vitamin B12 (0.1) | 256.8 | 8.326 | 240.5 to 273.1 |
| Zinc (50) | 256.4 | 32.63 | 192.4 to 320.3 |
| Vitamin K (10) | 252.4 | 53.2 | 148.2 to 356.7 |
| Vitamin E (0.01) | 248.6 | 17.08 | 215.1 to 282.0 |
| Vitamin B12 (500) | 247.6 | 26.48 | 195.7 to 299.5 |
| Copper (10) | 246 | 28.16 | 190.8 to 301.2 |
| Chromium (500) | 244.7 | 64.92 | 117.5 to 371.9 |
| Vitamin K (100) | 244 | 29.41 | 186.4 to 301.6 |
| Cobalt (100) | 242.5 | 22.6 | 198.2 to 286.8 |
| $\alpha$-linolenic acid (0.1) | 240.5 | 50.59 | 141.4 to 339.7 |
| Zinc (10) | 235.8 | 27.83 | 181.3 to 290.4 |
| Cobalt (50) | 233.2 | 16.94 | 200.0 to 266.4 |
| Copper (0.1) | 225.5 | 15.53 | 195.0 to 255.9 |
| Vitamin E (10) | 222.3 | 33.49 | 156.6 to 287.9 |
| Chromium (0.01) | 221 | 33.23 | 155.9 to 286.2 |
| Vitamin K (1) | 220.9 | 43 | 136.6 to 305.2 |

| | | | |
|---|---|---|---|
| Zinc (100) | 220.4 | 31.27 | 159.1 to 281.7 |
| Vitamin D (1) | 219 | 19.6 | 180.6 to 257.4 |
| Iodine (0.1) | 217.6 | 58.59 | 102.8 to 332.4 |
| Thyroxine (0.1) | 215.8 | 41.92 | 133.6 to 298.0 |
| BR + TGFβ1 | 215.4 | 25.49 | 165.4 to 265.3 |
| Vitamin D (0.1) | 210.1 | 12.26 | 186.1 to 234.2 |
| Zinc (1) | 207.1 | 15.94 | 175.9 to 238.4 |
| Vitamin D (10) | 204.7 | 24.25 | 157.1 to 252.2 |
| Vitamin A (0.01) | 203.5 | 30.72 | 143.3 to 263.7 |
| Zinc (0.1) | 202 | 28.98 | 145.2 to 258.8 |
| Manganese (0.1) | 200 | 36.97 | 127.5 to 272.4 |
| Manganese (50) | 199 | 28.23 | 143.7 to 254.4 |
| Vitamin E (1) | 198 | 11.83 | 174.8 to 221.1 |
| Cobalt (1) | 197.1 | 36.02 | 126.5 to 267.7 |
| Iodine (50) | 196.8 | 50.85 | 97.16 to 296.5 |
| α-linolenic acid (0.01) | 190.1 | 41.5 | 108.7 to 271.4 |
| Vitamin E (0.1) | 189.2 | 16.26 | 157.3 to 221.0 |
| Vitamin B7 (0.1) | 187.7 | 48.11 | 93.42 to 282.0 |
| α-linolenic acid (10) | 186.2 | 10.96 | 164.8 to 207.7 |
| Molybdenum (0.1) | 185.1 | 41.19 | 104.4 to 265.8 |
| α-linolenic acid (1) | 185.1 | 15.99 | 153.8 to 216.5 |
| Manganese (500) | 183.7 | 42.03 | 101.3 to 266.0 |
| Vitamin B7 (0.01) | 181.3 | 27.8 | 126.8 to 235.8 |
| Chromium (100) | 180.7 | 12.43 | 156.4 to 205.1 |
| Cobalt (0.1) | 180.4 | 17.91 | 145.3 to 215.5 |
| Vitamin E (50) | 177.2 | 3.812 | 169.7 to 184.7 |
| Manganese (0.01) | 172.9 | 15.97 | 141.6 to 204.2 |
| Copper (50) | 167.7 | 13.26 | 141.7 to 193.7 |
| Thyroxine (0.01) | 167.1 | 62.27 | 45.00 to 289.1 |
| Chromium (1) | 162.1 | 6.33 | 149.7 to 174.5 |
| Vitamin B7 (50) | 159.6 | 16.71 | 126.9 to 192.4 |
| Thyroxine (50) | 158.5 | 12.43 | 134.2 to 182.9 |
| Zinc (500) | 156.7 | 52.09 | 54.57 to 258.7 |
| Molybdenum (50) | 156.5 | 16.05 | 125.0 to 187.9 |
| Iodine (500) | 155.7 | 14.64 | 127.0 to 184.4 |
| Chromium (10) | 155.3 | 34.66 | 87.38 to 223.2 |
| Molybdenum (500) | 155 | 29.99 | 96.24 to 213.8 |
| Molybdenum (0.01) | 154.2 | 15.7 | 123.5 to 185.0 |
| Iodine (0.01) | 152.8 | 10.71 | 131.8 to 173.8 |
| Cobalt (500) | 152.7 | 12.46 | 128.3 to 177.1 |
| Manganese (1) | 152.2 | 22.58 | 108.0 to 196.5 |
| Thyroxine (1) | 149.6 | 13.24 | 123.6 to 175.5 |
| Vitamin B7 (1) | 147.5 | 15.61 | 116.9 to 178.1 |

| | | | |
|---|---|---|---|
| Manganese (10) | 144.3 | 13.18 | 118.5 to 170.2 |
| Vitamin B7 (500) | 143.1 | 7.943 | 127.5 to 158.6 |
| Thyroxine (10) | 141.2 | 7.162 | 127.2 to 155.2 |
| Iodine (10) | 139.5 | 8.329 | 123.2 to 155.8 |
| Vitamin D (50) | 139.1 | 23.62 | 92.79 to 185.4 |
| Molybdenum (1) | 136.5 | 13.72 | 109.6 to 163.4 |
| Iodine (100) | 135.2 | 6.792 | 121.9 to 148.5 |
| Manganese (100) | 132.5 | 12.14 | 108.7 to 156.3 |
| Vitamin B7 (10) | 125.7 | 9.282 | 107.5 to 143.9 |
| Vitamin B7 (100) | 124.6 | 14.03 | 97.15 to 152.2 |
| Molybdenum (10) | 124.4 | 16.84 | 91.35 to 157.4 |
| Molybdenum (100) | 120.1 | 4.995 | 110.3 to 129.9 |
| Iodine (1) | 119.6 | 6.093 | 107.7 to 131.5 |
| Thyroxine (100) | 117.3 | 12.23 | 93.30 to 141.2 |
| Thyroxine (500) | 116.2 | 11.47 | 93.67 to 138.6 |
| Vitamin E (500) | 113.8 | 13.05 | 88.21 to 139.4 |
| Vitamin D (100) | 104.3 | 10.17 | 84.41 to 124.3 |
| Vitamin D (500) | 56.04 | 1.154 | 53.78 to 58.30 |
| Copper (500) | 54.15 | 4.816 | 44.71 to 63.59 |
| Vitamin A (0.1) | 53.1 | 5.784 | 41.76 to 64.44 |
| Vitamin A (1) | 38.77 | 12.79 | 13.70 to 63.84 |
| $\alpha$-linolenic acid (100) | 38.66 | 1.248 | 36.22 to 41.11 |
| Vitamin A (100) | 38.49 | 15.45 | 8.204 to 68.78 |
| Vitamin A (10) | 36.95 | 7.876 | 21.51 to 52.38 |
| Vitamin A (50) | 16.57 | 1.479 | 13.67 to 19.46 |
| Vitamin A (500) | 12.13 | 0.9688 | 10.23 to 14.03 |
| $\alpha$-linolenic acid (500) | -14.43 | 0.5707 | -15.55 to -13.31 |

**Supplemental Data 7**

The effect of vitamins and minerals on *Gaussia* luciferase luminescence was assessed by supplementation of conditioned medium with each of the vitamins and minerals tested. No significant changes were seen.

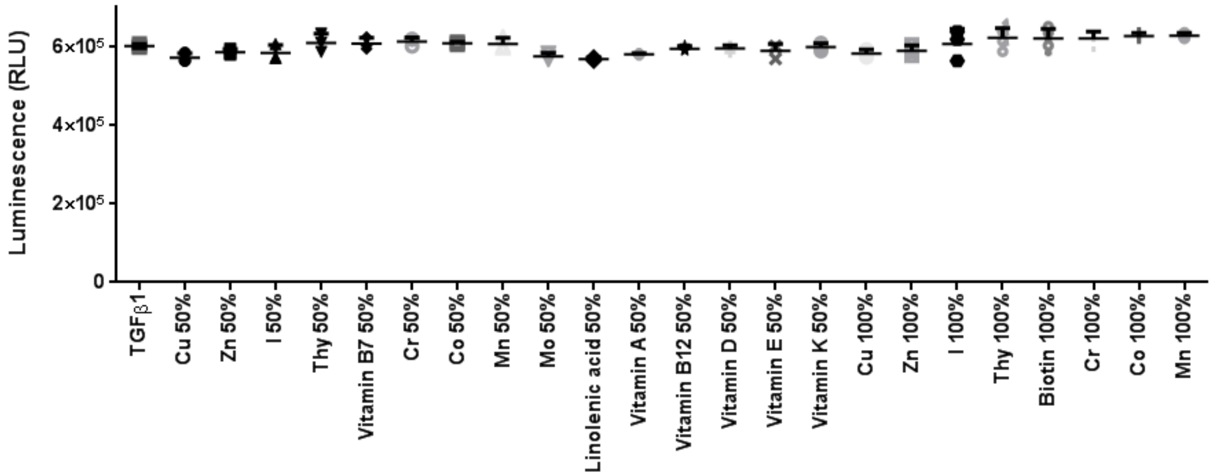
